# Inhibition of retrotransposition improves health and extends lifespan of SIRT6 knockout mice

**DOI:** 10.1101/460808

**Authors:** Matthew Simon, Michael Van Meter, Julia Ablaeva, Zhonghe Ke, Raul S. Gonzalez, Taketo Taguchi, Marco De Cecco, Katerina I. Leonova, Valeria Kogan, Stephen L. Helfand, Nicola Neretti, Asael Roichman, Haim Y. Cohen, Marina Antoch, Andrei Gudkov, John M. Sedivy, Andrei Seluanov, Vera Gorbunova

## Abstract

Mice deficient for SIRT6 exhibit a severely shortened lifespan, growth retardation, and highly elevated LINE1 (L1) activity. Here we report that SIRT6 deficient cells and tissues accumulate abundant cytoplasmic L1 cDNA which triggers massive type I interferon response via activation of cGAS. Remarkably, nucleoside reverse transcriptase inhibitors (NRTIs), which inhibit L1 retrotransposition, significantly improved health and lifespan of SIRT6 knockout mice and completely rescued type I interferon response. In tissue culture, inhibition of L1 with siRNA or NRTIs abrogated type I interferon response, in addition to a significant reduction of DNA damage markers. These results indicate that L1 activation contributes to the pathologies of SIRT6 knockout mice. Similarly, L1 transcription, cytoplasmic cDNA copy number and type I interferons were elevated in the wild type aged mice. As sterile inflammation is a hallmark of aging we propose that modulating L1 activity may be an important strategy for attenuating age-related pathologies.

**Highlights:**

- SIRT6 KO mice accumulate L1 cDNA triggering type I interferon response via cGAS pathway
- Wild type aged mice accumulate L1 cDNA and display type I interferon response
- Reverse transcriptase inhibitors rescue type I interferon response and DNA damage
- Reverse transcriptase inhibitors extend lifespan and improve health of SIRT6 KO mice

## INTRODUCTION

The process of progressive organismal and cellular decline known as aging remains one of the most challenging problems in biology and medicine, resulting as the ultimate cause of morbidity and mortality. Investigating and understanding the underlying mechanisms that promote aging is of paramount importance to elucidating interventions in age-related diseases. Of the multiple observations that have been made in aging systems, one of the more mysterious has been the role of the transposable elements that have colonized the mammalian genome. While most of these elements lack the ability to replicate beyond their host cell, they are nonetheless still capable of replicating their DNA and pose a potential threat to the integrity of the host genome. One of the more successful retrotransposons is the class of long interspersed nuclear element 1 (L1), which is a ubiquitous feature of mammalian genomes, comprising approximately 20% of the genomic DNA in mice and humans (Lander et al., 2001; Mouse Genome Sequencing et al., 2002). These 6 kb, fully functional retrotransposons can not only replicate themselves, but also other retroelements that use L1-encoded proteins necessary for retrotransposition: ORF1, a nucleic acid chaperone, and ORF2, an endonuclease and reverse transcriptase (Dewannieux et al., 2003; Hancks and Kazazian, 2012; Richardson et al., 2015). As this ability to replicate and expand within a host genome requires DNA breakage and insertion, L1 activity has been linked to DNA damage and mutagenesis (Gasior et al., 2006; Gilbert et al., 2002; Iskow et al., 2010).

While the preponderance of research on L1s has focused on their activity in the germ line, recent evidence has suggested that L1 activity in somatic tissues contributes to a number of age-related diseases, such as neurodegeneration and cancer (Hancks and Kazazian, 2012; Iskow et al., 2010; Lee et al., 2012; Reilly et al., 2013; Tan et al., 2012). Given the potential harm that L1s can cause, host genomes have evolved a number of molecular mechanisms for silencing these parasitic elements (Crichton et al., 2014; Levin and Moran, 2011). Recent studies have reported, however, that these mechanisms become less efficient during the aging process, resulting in the derepression of L1s (De Cecco et al., 2013a; St Laurent et al., 2010; Van Meter et al., 2014). Much of this derepression appears to stem from redistribution and reorganization of the heterochromatin that normally constrains the activity of these elements and can lead to inflammation through the innate immune response (Ablasser et al., 2014; Oberdoerffer et al., 2008; Thomas et al., 2017; Van Meter et al., 2014). Despite the results which suggest that upregulation of L1 expression is a hallmark of aging, it is unclear to what extent, if any, L1s contribute to age-related pathologies and whether inhibiting L1 activity can delay these pathologies.

Mice deficient in the mono-ADP-ribosylase/deacetylase protein SIRT6 develop a severe premature aging phenotype, characterized by a failure to thrive, intestinal sloughing, hypoglycemia and a severely shortened lifespan (Mostoslavsky et al., 2006). Recently, SIRT6, has been demonstrated to be involved in silencing of L1 promoters (Van Meter et al., 2014). SIRT6 mono-ADP-ribosylates KAP1 and promotes its complex formation with HP1 thereby packaging the L1 DNA into transcriptionally silent heterochromatin (Van Meter et al., 2014). Remarkably, SIRT6 knockout (KO) mice show a strong activation of L1, suggesting a role for L1 misexpression in their age-related phenotypes (Mostoslavsky et al., 2006; Van Meter et al., 2014). Additionally, SIRT6 KO cells display excess genomic instability and DNA damage (Mostoslavsky et al., 2006). Considering the short lifespan of SIRT6 deficient mice, they provide a unique model to test whether inhibition of L1 activity can extend lifespan and alleviate the pathology of these mice.

Given the multiple progeroid phenotypes and highly elevated L1 activity inherent to SIRT6 KO mice, we sought to use this system to address the role for L1s in age-related pathology. We used anti-reverse transcriptase drugs to treat SIRT6 KO cell lines and animals to inhibit L1 retrotranspositon and found that many of the pathologies in these animals were alleviated. L1-specific RNAi knockdown was able to recapitulate the results with the anti-reverse transcriptase drugs. We also discovered that SIRT6 KO associated L1 activity triggers activation of the innate immune response via type I interferon production. Finally, we show that L1 DNA accumulates in the cytoplasm in SIRT6 KO tissues, triggering the cytoplasmic DNA sensor, cGAS, and initiating the innate immune response. Our data reveal that L1 activity directly contributes to the progeroid phenotypes of SIRT6 mice and correlates with observations in normal aging tissues and animals.

## RESULTS

### NRTI treatment abrogates L1 activity in SIRT KO cells and tissues

Nucleoside reverse transcriptase inhibitors (NRTI) are a powerful class of clinical antiviral compounds used to treat HIV-1 infection via poisoning of reverse transcriptase (RT) enzymes by terminating chain elongation (Painter et al., 2004). This activity effectively blocks retroviruses and retroelements from completing genomic invasion. Several studies have reported that in addition to impeding the activity of viral polymerases, NRTIs such as lamivudine and stavudine, are also potent inhibitors of L1 RT (Dai et al., 2011; Jones et al., 2008; Xie et al., 2011). In order to assess the efficacy of NRTIs in inhibiting L1 in the context of SIRT6 KO, we transfected WT and SIRT6 KO mouse embryonic fibroblasts with either human or mouse L1-enhanced green fluorescent protein (EGFP) reporter cassettes (Ostertag et al., 2000) to measure *de novo* retrotransposition events. In brief, successful retrotranspositon is detected when the GFP marker is retrotranscribed with the interrupting intron spliced out during mRNA processing. Successful events are measured as percent of GFP-positive cells. L1s were approximately 3- times more active in SIRT6 KO relative to WT cells (**Figure 1A, B**; **Figure S1A**). Both stavudine or lamivudine treatments abrogated L1 retrotransposition events in both WT and SIRT6 KO cells, demonstrating a robust antagonistic activity to the L1 lifecycle (**Figure 1A, B**; **Figure S1A**). Additionally, we found that SIRT6 KO MEFs demonstrate a progressive accumulation of L1 DNA with each population doubling (PD) (**Figure 1C**). NRTI treatment of WT and SIRT6 KO fibroblasts over the course of 40 PDs effectively inhibited the expansion of L1 DNA copies in SIRT6 KO cells, demonstrating that NRTIs are sufficient for ameliorating L1 DNA accumulation (**Figure 1C**).

**Figure 1.**
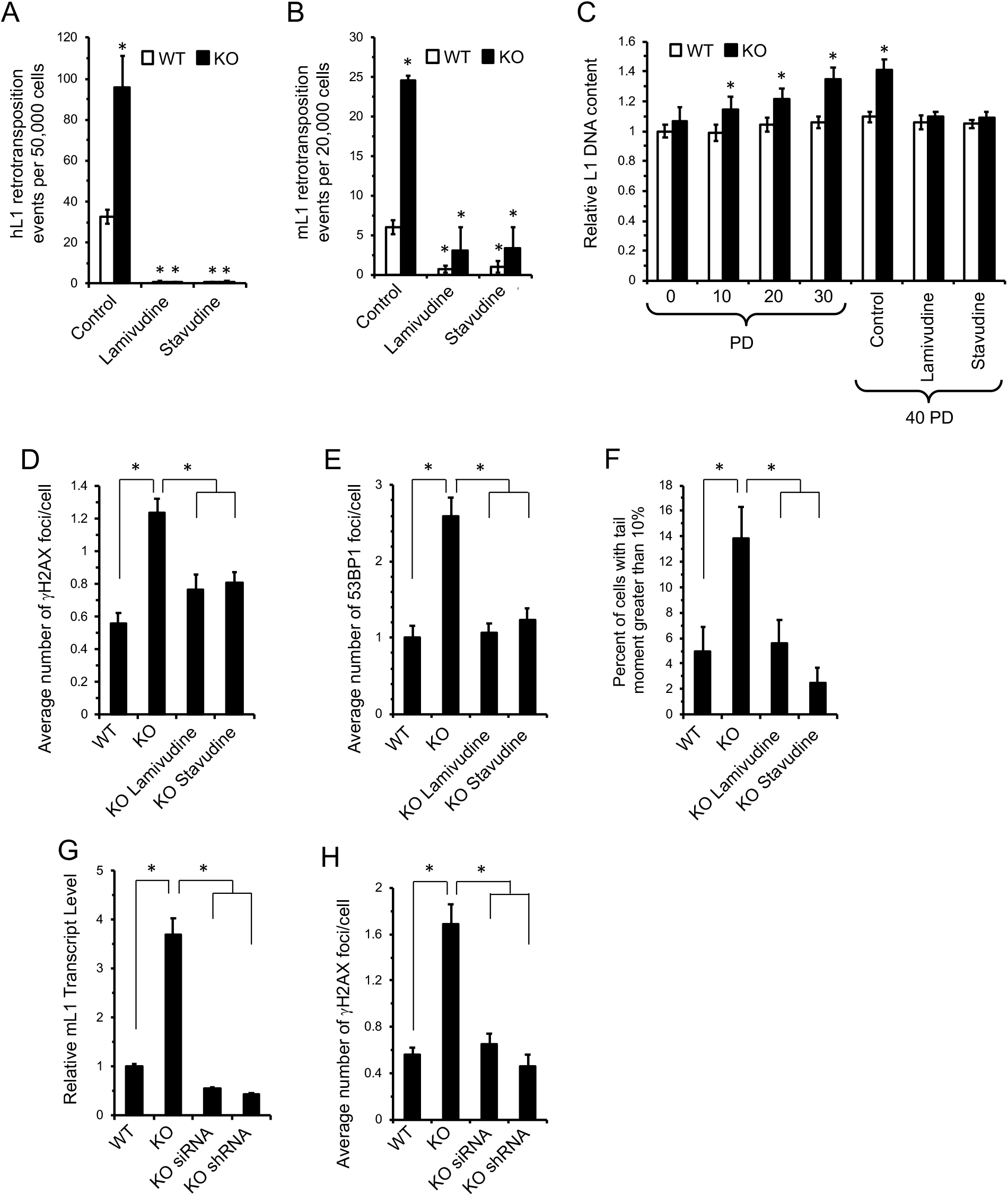
NRTI treatment inhibits L1 retrotransposition and rescues DNA damage in SIRT6 KO cells. **A, B** Treatment with either stavudine or lamivudine inhibits L1 retrotransposition events. WT and SIRT6 KO MEF were cultured with 10 μM stavudine or lamivudine and then transfected with a human (**A**) or mouse (**B**) L1-EGFP reporter plasmid^21^. *De novo* retrotransposition events leading to activation of the GFP gene were scored by flow cytometry. The values were normalized to actin. **C**, NRTI treatment reduces L1 DNA copy number in cultured cells. WT and SIRT6 KO MEFs were cultured and assayed for L1 copy number every 10 population doublings (PDs). In addition (right part of the panel), cells were grown for 40 PDs with 10 μM NRTI and then assayed by qPCR. The values were normalized to 5S ribosomal RNA gene. **D-F**, SIRT6 KO MEFs show elevated levels of DNA damage that is alleviated by NRTI treatment. MEFs were isolated from embryos of NRTI-treated or control dams, cultured for two passages with or without NRTIs and spontaneously arising γH2AX (**D**) and 53BP1 (**E**) foci were quantified by immunostaining. At least 80 cells were counted for each treatment. **F**, SIRT6 KO MEFs show elevated levels of DSBs as measured by the neutral comet assay, and these breaks are rescued by NRTI treatment. Values represent percent of population with excessive DNA damage denoted by tail DNA content in excess of 10%. At least 80 cells were counted for each treatment. **G**, L1-specific knockdown rescues elevated L1 expression in SIRT6 KO cells. SIRT6 KO MEF lines were generated with siRNA or shRNA cassettes targeting conserved sequences of L1. The cassettes stably integrated. Both cassettes rescued L1 misexpression. qRT-PCR data was normalized to actin. **H**, L1 RNAi rescued elevated γH2AX foci in SIRT6 KO cell lines. For qRT-PCR experiments, three independent MEF cultures of each genotype were used. Error bars show s.d. Statistical significance was determined by the Student’s *t*-test and asterisks indicate *p* < 0.05.

### Inhibition of L1s rescues elevated DNA damage in SIRT6 KO cells

DNA breaks induced by L1 ORF2 protein and *de novo* insertion events pose a threat to genomic stability. Overexpression of ORF2p is known to induce excessive DNA damage and can induce senescence, demonstrating the potential danger in endogenous misregulation of L1 activity (Gasior et al., 2006; Gilbert et al., 2002; Kines et al., 2014). In order to assess the effect of NRTI treatment on L1-mediated DNA damage, we treated mouse fibroblasts with NRTIs for 10 PDs. SIRT6 KO cells showed elevated double strand breaks by both γH2AX (**Figure 1D**, **Figure S1B**) and 53BP1 staining (**Figure 1E**, **Figure S1B**) and by neutral comet assay (**Figure 1F**). This DNA damage was significantly ameliorated by NRTI treatment (**Figure 1D-F**, **Figure S1B**). Thus, NRTI treatment significantly reduces genomic instability linked to L1 RT in SIRT6 KO cells.

NRTIs function as broad reverse transcriptase inhibitors. SIRT6 KO tissues demonstrate misregulation and increased expression of many retroelements, including SINEs and other retrotransposons (Van Meter et al., 2014), any of which could also contribute to SIRT6 KO phenotypes and would potentially be suppressed by NRTI treatment. To elucidate the direct role of L1 activity in SIRT6 KO biology, we generated two L1 RNAi vector systems to directly target L1s using conserved sequences between different L1 families, especially the more active families, including L1MdA_I, II, III and L1MdTf (Hardies et al., 2000; Sookdeo et al., 2013). RNAi cassettes, or control vectors, were integrated into SIRT6 KO MEFs. Both RNAi systems demonstrated significant reduction in L1 RNA abundance (**Figure 1G**). Immunostaining was conducted on these cell lines to assess DNA damage. Strikingly, both RNAi systems rescued the elevated γH2AX foci observed in the SIRT6 KO cells (**Figure 1H**). Thus, L1 activity alone in SIRT6 KO cells is the major contributor to the excessive DNA damage observed.

### Elevated L1 activity in SIRT6 KO mouse organs is suppressed by NRTI treatment

Based on the successful *in vitro* inhibition of L1 RT activity in SIRT6 KO cells and the significant amelioration of cellular DNA damage, we began administering stavudine and lamivudine to SIRT6 KO mice. Heterozygous SIRT6^+/-^ mice were bred and pregnant animals were administered either stavudine or lamivudine in the drinking water immediately after mating. NRTIs continued to be administered in the drinking water throughout the postnatal period; in addition, the pups were given NRTIs orally once a day using a pipette, while the control pups were given water. As expected, in the homozygous SIRT6 KO control group, multiple tissues showed upregulation of L1 transcription compared to WT littermates (**Figure 2A**). Total L1 DNA content (**Figure 2B**) was also highly elevated in these tissues. Remarkably, this increase in L1 DNA was suppressed by NRTI treatments (**Figure 2B**). Additionally, WT littermate cohorts treated with NRTIs also demonstrated significant decreases in L1 DNA content (**Figure 2C**). These data demonstrate that NRTIs can suppress L1 activity to a significant extent in these animals, regardless of the baseline level of L1 activity.

**Figure 2.**
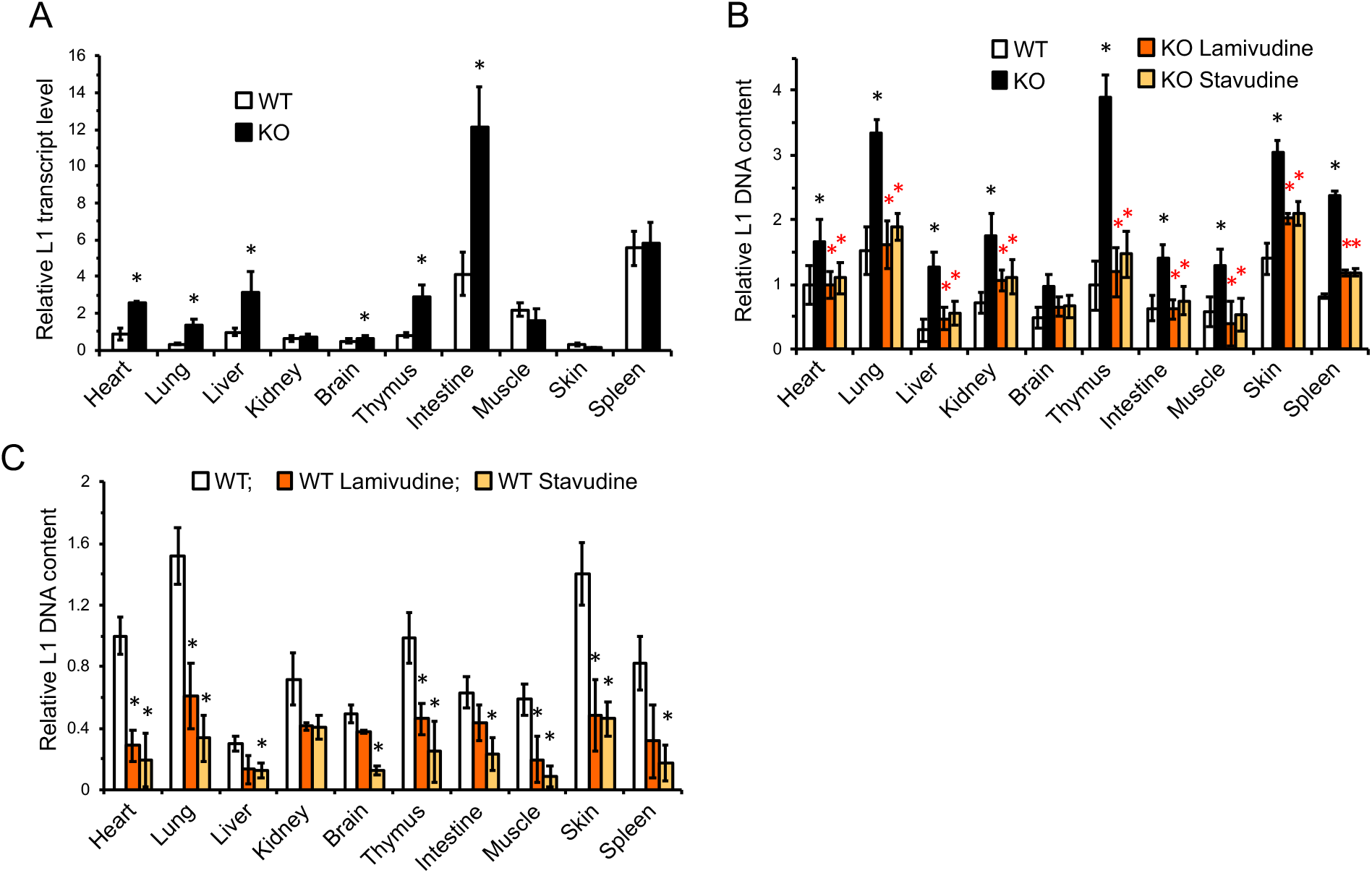
NRTIs suppress L1 activity in WT and SIRT6 KO mice. **A,** L1 transcripts are elevated in several tissues of SIRT6 KO mice. Organs were harvested from 27 days old WT and SIRT6 KO mice. L1 mRNA was assayed via qRT-PCR and normalized to actin and WT heart was used as a reference. **B**, NRTI treatment reduces L1 DNA content in SIRT6 KO mice. **C,** L1 DNA content in NRTI treated WT mouse tissues. Mice were treated with NRTIs as described in Methods. Tissues were assayed for L1 DNA content by qPCR at 27 days old. The values were normalized to 5S rRNA gene. Four animals were assayed for each treatment and error bars indicate s.d. Statistical significance was determined by the Student’s *t*-test and asterisks indicate *p* < 0.05.

### Cytoplasmic L1 DNA is enriched in SIRT6 KO cells and tissues

Previous studies reported that conditions such as autoimmunity (Thomas et al., 2017), (Stetson et al., 2008), cancer (Shen et al., 2015) or HIV infection (Jones et al., 2013) are associated accumulation of extrachromosomal L1 cDNA copies. Additionally, it has been reported that extranuclear DNA sensing is essential for cellular senescence (Dou et al., 2017; Li and Chen, 2018; Takahashi et al., 2018; Yang et al., 2017), which is elevated with SIRT6 deficiency and increases in aging mammals (Mao et al., 2012; Nagai et al., 2015). To test whether an increased L1 copy number is associated with the increase in the extrachromosomal L1 cDNA copies, immunofluorescence staining was performed using an anti-ssDNA antibody. Extranuclear ssDNA foci were consistently observed in SIRT6 KO cells, but not in WT cells (**Figure 3A**). Further, FISH staining using a L1 DNA-specific probe revealed multiple foci in SIRT6 KO cells, indicating that L1 DNA is present to a significant degree in the cytoplasm of these cells. Cell fractionation and subsequent qPCR quantification demonstrated SIRT6 KO fibroblasts contained 2-fold greater number of cytoplasmic L1 DNA than the WT cells, and this increase was completely rescued by the NRTIs (**Figure 3B**). Similarly, cytoplasmic L1 DNA content was also highly elevated in tissues that had demonstrated high L1 expression and total DNA content (**Figure 3C-F**). These data indicate that much of the elevated L1 DNA observed in SIRT6-deficient tissues are not integrated copies and exist as extra-chromosomal cytoplasmic DNA.

**Figure 3.**
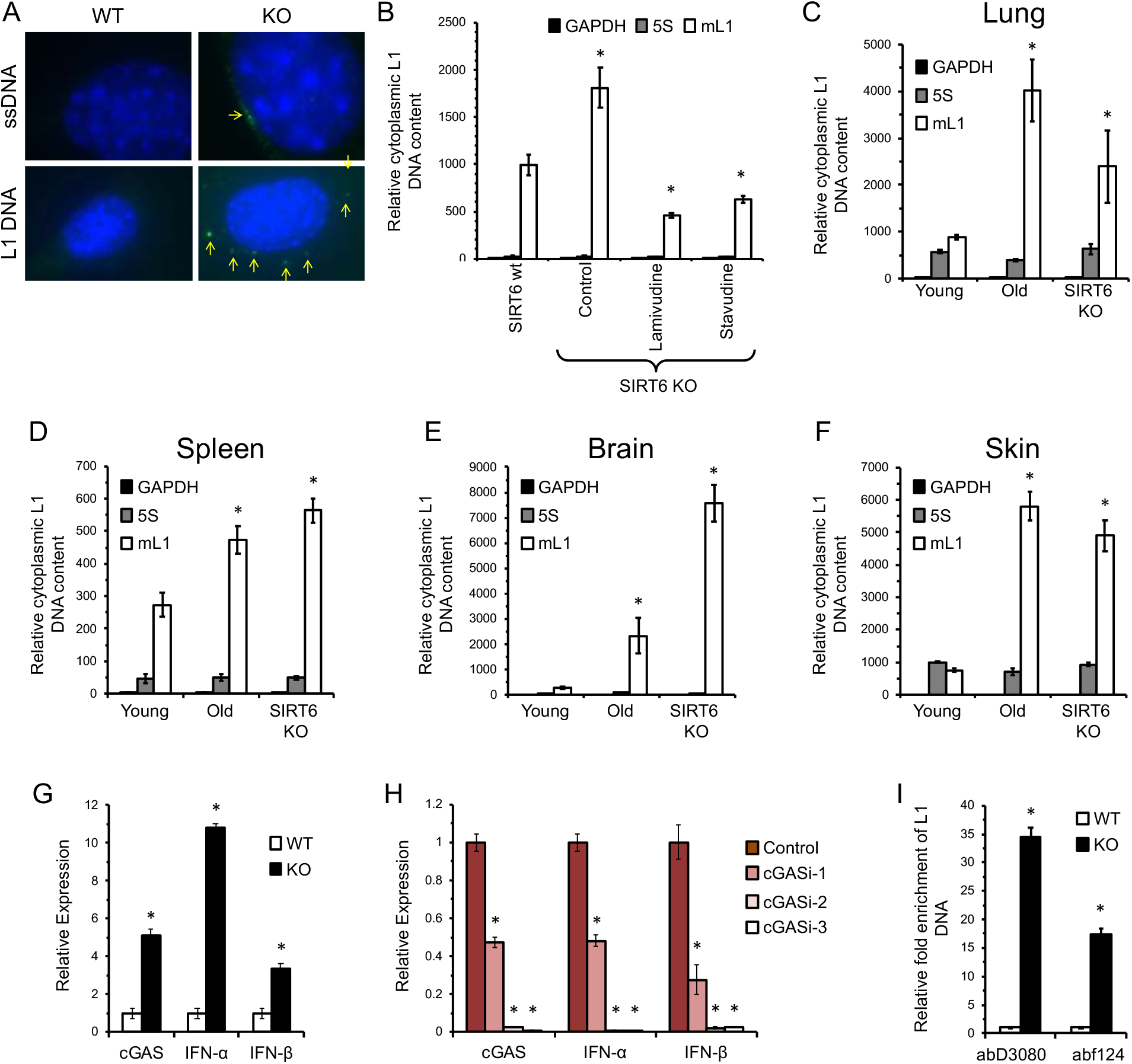
Cytoplasmic L1 DNA is enriched in SIRT6 KO and aged WT tissues. **A**, SIRT6 KO cells have cytoplasmic ssDNA and L1 DNA. Cytoplasmic ssDNA foci were observed in cultured SIRT6 KO MEFs but were not observed in WT cells. Similarly, SIRT6 KO MEFs staining using a DNA FISH probe displayed multiple foci in the cytoplasm that were rare or absent in WT cells. **B,** Cytoplasimic L1 DNA is elevated in SIRT6 KO MEFs. Cells cultured with NRTIs show reduced cytoplasmic L1 DNA content. Cells were counted and lysed to extract cytoplasmic fraction. GAPDH and ribosomal subunit 5S primers were used to detect nuclear contamination and did not show statistically significant differences between treatments. **C-F**, Cytoplasmic L1 DNA copies are elevated in the tissues of aged (24 month old) and SIRT6 KO mice. **C**, liver; **D**, spleen; **E**, brain; **F**, skin. **G**, cGAS expression and type I interferon expression are elevated in SIRT6 KO MEFs. Expression levels were normalized to actin. **H**, RNAi knockdown of cGAS correlates with comparable decreases in type I interferon expression in SIRT6 KO MEFs. 3 separate cGAS targeting shRNAs were transfected into MEF SIRT6 KO cells. A scramble shRNA sequence was used as a control. Expression levels were normalized to actin. **I,** cGAS-bound DNA is enriched for L1s in SIRT6 KO cells. Immunoprecipitation of cGAS from MEFs using two separate antibodies shows ~17 and ~34-fold enrichment for L1 DNA. Abundance of cGAS-bound DNA was normalized to 5S ribosomal subunit DNA. Three independent MEF cultures or four animals of each genotype were used. Error bars show s.d. Statistical significance was determined by the Student’s *t*-test and asterisks indicate *p* < 0.05.

### L1 activity correlates with type I interferon response and is rescued by NRTI treatment

L1 transposition intermediates such as L1 RT reverse transcribed cytoplasmic cDNA can be recognized by cellular antiviral defense machineries triggering a type I interferon response (Volkman and Stetson, 2014). Indeed, we observed a significant increase in total L1 DNA copies, as well as cytoplasmic L1 cDNA in SIRT6 KO cells and tissues (**Figure 3B-F**) which could trigger a type I interferon response (McNab et al., 2015). In SIRT6 KO MEFs, the cytosolic DNA sensor, cGAS, exhibited significantly higher expression, which coincided with elevated expression of type I interferons (IFN-α and IFN-β1) (**Figure 3G**). To confirm that cGAS signaling was responsible for the observed interferon response, RNAi against cGAS was used. Using 3 separate shRNAs with varying degrees of efficacy, cGAS knockdown correlated with suppression of both IFN-α and IFN-β1 (**Figure** 3H). Finally, MEF cells were crosslinked with UV radiation and cGAS was immunoprecipitated using two separate antibodies. Subsequent purification and analysis of the bound DNA revealed a ~17 and ~34 –fold increase in the abundance of L1 DNA in SIRT6 KO cells compared to WT, depending of the antibody used (**Figure 3I**). Thus, cytosolic L1 DNA triggers type I interferon expression via the cGAS signaling pathway.

Consistent with the results in MEFs, multiple tissues of SIRT6 KO mice showed a dramatic increase in type I interferon (IFN-α and IFN-β1) expression, which was completely rescued by NRTI treatment (**Figure 4A, B**). Additionally, both L1 RNAi systems completely rescued IFN-α and IFN-β1 expression, demonstrating similar trends to those observed in the NRTI-treated animals and indicating that L1 activity is the cause of the innate immune response (**Figure 4C, D**). Further, suppression of basal levels of type I interferons by NRTI treatment was also observed in WT tissues (**Figure S2A, B**). Other inflammation markers were also elevated in SIRT6 KO mice, many of which were reversed by NRTI treatment (**Figure S2C**).

**Figure 4.**
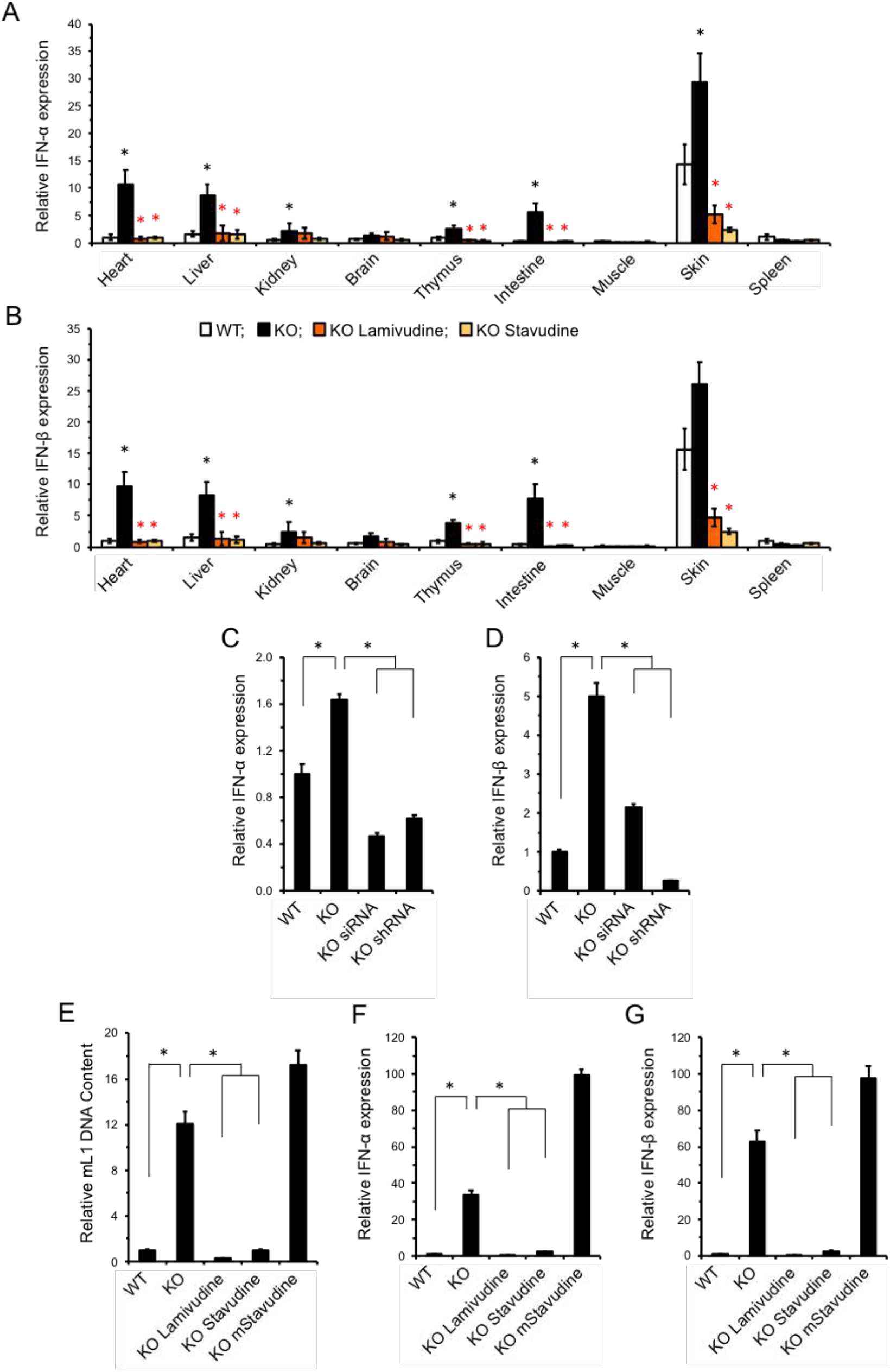
Inhibition of reverse transcriptase rescues type I interferon response in SIRT6 KO mice. **A, B**, SIRT6 KO mice display a robust type I interferon-α (**A**) and β (**B**) response that is rescued by NRTI treatment. Organs were harvested from 27 days old mice and interferon mRNA was quantified by qRT-PCR. The data was normalized to actin and WT heart was used as a reference. Five animals were assayed for each group and error bars show s.d. Black asterisks indicate significant differences from WT, and red asterisks indicate significant differences from water-treated SIRT6 KO. **C, D,** SIRT6 KO MEF lines were generated with stably integrated siRNA or shRNA cassettes targeting conserved sequences of L1. These cassettes efficiently repress L1 transcription (Figure 1G. Both cassettes rescued IFN-α/β expression (**C, D**) **E-G**, NRTI anti-RT activity is essential for L1 suppression and rescue of type I interferon response. Cells were treated with 10 μM/ml of unmodified NRTI or methylated stavudine for 30 PD. L1 DNA content (**E**) and IFN-α/β expression (**F**, **G**) were assayed by qPCR and qRT-PCR. The data was normalized to actin and WT values were used as a reference. Statistical significance was determined by the Student’s *t*-test and asterisks indicate values significantly different from the wild type (*p* < 0.05).

Previously, it was reported that that NRTIs possess an intrinsic anti-inflammatory activity independent of their anti-RT activity (Fowler et al., 2014), raising the possibility that the observed rescue of interferon response is independent of L1 inhibition. To address this possibility, we synthesized a 5’-O-methyl (meStav) version of stavudine, which was previously reported to lack anti-RT activity due to the removal of the 5’-OH group but still retained anti-inflammatory activity (Fowler et al., 2014). SIRT6 KO MEFs treated with meStav at the same dosage as stavudine failed to suppress IFN-α and IFN-β1 expression, indicating that NRTI anti-RT activity is essential for suppression of the innate immune response (**Figure 4E-G**).

### NRTI-treatment alleviates progeroid phenotypes and extends lifespan in SIRT6 KO mice

Consistent with previous studies (Mostoslavsky et al., 2006), the control (water treated) SIRT6 KO mice rapidly developed progeria and postnatal wasting with complete penetrance. Remarkably, the NRTI-treated mice presented as generally healthier, with shiny fur, improved body size, and less kyphosis (**Figure 5A**). While the control SIRT6 KO mice all died within 35 days, the NRTI-treated SIRT6 KO mice exhibited a more than two-fold increase in their mean and maximum lifespans (**Figure 5B**). Additionally, we observed that NRTI-treated SIRT6 KO mice had improved body mass and delayed wasting compared to the controls (**Figure 5C**). Thus, NRTI treatment significantly improves SIRT6 KO lifespan and healthspan.

**Figure 5.**
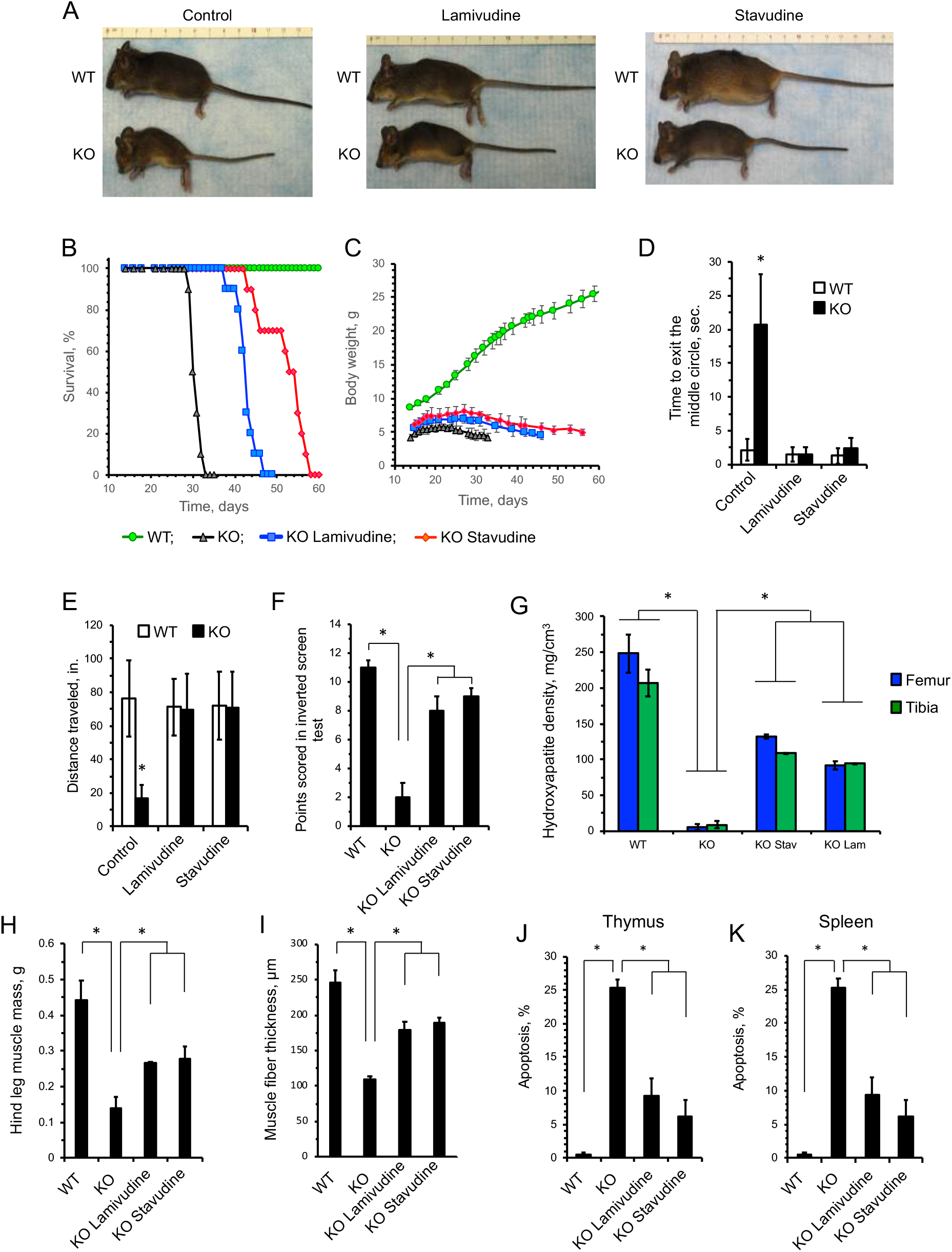
Treatment with NRTIs extends lifespan and improves health of SIRT6 KO mice. Mice were treated as described in Methods. **A,** NRTI treatment improves the appearance and alleviates wasting phenotype of SIRT6 KO animals. Pictures were taken at 30 days of age. **B,** NRTI treatment extends mean and maximum lifespans of SIRT6 KO mice. Each treatment group contained 10 animals. The survival curves for SIRT6 KO NRTI-treated mice were significantly different from untreated SIRT6 KO controls. *p* < 0.0001, log-rank test. **C,** NRTI treatment improved body weight of SIRT6 KO animals. Each treatment group contained 10 animals. *p* < 0.0001, One Way ANOVA. **D-F,** NRTI treatment rescues physical impairments in SIRT6 KO mice. **D,** Open field test: healthy mice avoid unfamiliar open spaces and rapidly run to the corners when placed in the center. Time to exit the field was measured. **E,** Foraging activity. Animal movements were recorded for 30 seconds using a camera, and the distance traveled was quantified using Tracker software. **F,** Inverted screen test measures the time a mouse holds on to an inverted mesh screen. For each animal the point score was calculated as a sum of the scores for three trials, then an average was calculated for six animals. Six animals were assayed for each group, and three replicate tests were performed for each animal. **G,** Bone density of SIRT6 KO animals is significantly reduced, and is partially rescued by NRTI treatment. Data were determined by CT-scan and analysis of 27-day old animals. Three animals were assayed for each group and error bars show s.d. **H, I,** Hind leg muscle mass (**H**) and fiber thickness (**I**) are significantly reduced in SIRT6 KO animals and are partially rescued by NRTI treatment. Hind quarters (feet, tibia, femur were separated at pelvis, including all muscle groups, with exception of gluteus maximous) were removed and weighed. Fiber thickness was measured in quadriceps muscles fixed and stained with hematoxylin & eosin. At least 50 fibers were measured for cross sectional diameter per group at similar lateral locations on the quadriceps. Three animals were assayed for each group and error bars show s.d. **J, K**, NRTIs attenuate apoptosis in the thymus (**J**) and spleen (**K**) of SIRT6 KO mice. Apoptosis was measured in single cell suspensions by annexin V/PI staining and flow cytometry. Five mice were used for each group and error bars show s.d. (*p* < 0.01). Unless stated otherwise, statistical significance was determined by the Student’s *t*-test and asterisks indicate *p* < 0.05.

One of the phenotypes of SIRT6 KO mice is hypoglycemia (Xiao et al., 2010; Zhong et al., 2010), which was not rescued by NRTI treatment (**Figure S3A**). Blood glucose levels continued to decline with age in NRTI-treated SIRT6 KO animals, similar to the control animals (**Figure S3A**). Previous reports have indicated that SIRT6 KO postnatal wasting can be partially rescued by providing animals with supplemental glucose (Xiao et al., 2010). To test the potential contribution of blood glucose to NRTI-mediated rescue, we provided SIRT6 KO animals with 10% glucose with and without NRTIs. Glucose supplementation alone did not significantly improve median SIRT6 KO lifespan, but yielded two (out of ten) “survivor” mice that outlived the untreated KO mice (**Figure S3B**). However, when glucose supplementation was combined with the NRTIs it abrogated the NRTI rescue (**Figure S3C, D**). The fact that hypoglycemia was not rescued by NRTIs, and the negative impact of glucose supplementation on the lifespan of NRTI-treated animals indicates that NRTI-mediated rescue occurs independently of blood glucose changes.

In addition to improved body weight and lifespan, NRTI-treated animals showed dramatic improvements in mobility and behavior. Typically, SIRT6 KO animals demonstrate low mobility and unresponsiveness. However, NRTI-treated SIRT6 KO animals displayed a normal flight reflex from exposed locations (**Figure 5D** and Supplemental videos), foraging activity (**Figure 5E**), and greatly improved physical function on an inverted screen test (**Figure 5F**). Postnatal ophthalmological development is also stunted in SIRT6 KO mice, with pups unable to fully-open their eyes. NRTI-treated animals showed significant improvements over control animals (**Figure S4A-C**). SIRT6 KO animals demonstrated reduced bone density (**Figure 5G and Figure S4D**) and muscle mass and muscle fiber thickness (**Figure 5H, I and Figure S4E**), which were significantly improved by NRTI treatment. Additionally, SIRT6 KO animals showed deficiencies in hematopoietic (**Figure S5**) and intestinal (**Figure S6A**) stem cell compartments, which were significantly ameliorated by NRTI treatment. SIRT6 KO mice have also been reported to display severe lymphopenia (Mostoslavsky et al., 2006). In agreement with this, we observed elevated levels of apoptosis in thymus and spleen of the SIRT6 KO mice that were attenuated by the NRTI treatment (**Figure 5J, K**). Cumulatively, these data show that NRTI treatment drastically improves the healthspan of animals displaying elevated L1 activity, indicating an active role for L1s in these pathologies.

### NRTI treatment reduces apoptosis and improves health of SIRT6 KO intestines

SIRT6 KO mice have been previously reported to suffer from a severe colitis phenotype (Mostoslavsky et al., 2006). Intestines of SIRT6 KO animals exhibited dramatically reduced thickness of the epithelial layer, atrophied villi and pockets of inflammation, whereas the WT intestines did not (**Figure 6A, B**). Intestines of SIRT6 KO animals treated with either lamivudine or stavudine showed improved epithelial thickness and reduced inflammation (**Figure 6B, D**). In the small intestine, knockout mice displayed shortened villi (**Figure 6A, E**), decreased number of lamina propria plasma cells and lymphocytes, and pockets of neutrophilic acute inflammation dispersed throughout the lamina propria (**Figure 6B**). Additionally, apoptosis was significantly elevated in SIRT6 KO intestinal tissue and mucus-producing goblet cells were severely depleted (**Figure 6C, F, G**). NRTI treatment partially rescued these phenotypes, increasing villi size, vastly reducing debris, ameliorating apoptosis and suppressing inflammation (**Figure 6**). Restoration of tissue integrity in the intestine stands out in the context of extending SIRT6 KO lifespan, as the mice are characterized by a failure to thrive. Disruption of intestinal integrity is a well-characterized symptom of aging and perturbing intestinal permeability and barrier function is thought to contribute to age-related enteric diseases. No obvious histological differences were seen in the brains, kidneys, hearts, livers and lungs of the different mouse groups (**Figure S6B**).

**Figure 6.**
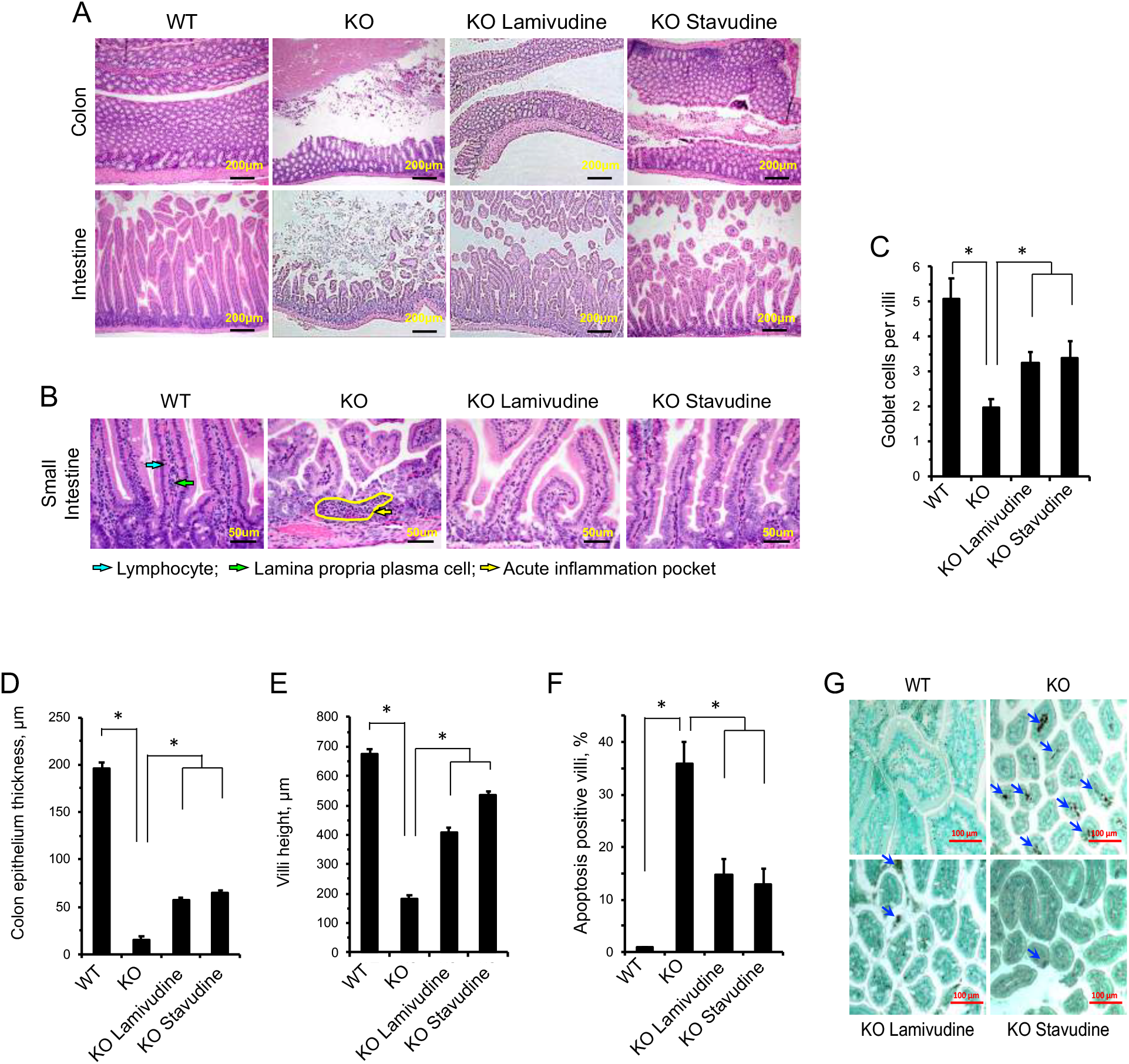
NRTIs improve gut health and reduce villi apoptosis in SIRT6 KO mice. NRTI treatment alleviates tissue pathology in the SIRT6 KO colon and small intestine. Histological examination was performed on four 27 day old animals for each group and representative images are shown. **A,** Tissues were stained with hematoxylin & eosin and are shown at 100x magnification (scale bars = 200 μm). **B,** Intestinal and colon tissues from WT and SIRT6 KO animals. SIRT6 KO small intestines displayed neutrophilic acute inflammation pockets (yellow arrow) and reduced villi size, while intestines from SIRT6 KO animals have reduced epithelial thickness. Normally abundant plasma cells (green arrow) and lymphocytes (blue arrow) found in WT tissues were rarely seen in SIRT6 KO lamina propria and were not restored by NRTI treatment. Tissues were stained with hematoxylin & eosin (scale bars = 50 μm). **C,** SIRT6 KO villi have depleted goblet cell populations that are partially restored by NRTI treatment. At least 20 villi were scored. **D, E**, NRTI treatment restores the thickness of the epithelial layer (**D**) and villi height (**E**) in SIRT6 KO intestines. **F, G**, NRTI treatment partially rescues elevated apoptosis observed in SIRT6 KO villi **(F)**. Apoptosis in the small intestine was analyzed by TUNEL staining. **G,** Representative images of TUNEL results in villi. Arrows indicate apoptotic cells. Statistical significance was determined by the Student’s *t*-test and asterisks indicate *p* < 0.05.

### L1 misexpression and sterile inflammation in aged mice coincide with cytoplasmic L1 DNA

Since sterile inflammation is a hallmark of aging (Lopez-Otin et al., 2013), we hypothesize that a type I interferon response mediated by age-related activation of L1 elements could play a causal role in this process. Consistent with this hypothesis, L1 expression and type I interferon expression were elevated in several tissues in the WT aged mice (**Figure 7A-C**), suggesting that L1 activation may be contributing to sterile inflammation associated with normal aging. Aged mice also showed increased L1 cytoplasmic DNA (**Figure 3C-F**). This result indicates that L1 activation is not limited to progeroid SIRT6 mice but is a hallmark of aging. Increase in L1 RNA and DNA in old tissues has been previously reported (De Cecco et al., 2013b). In order to assess if reverse transcription inhibition impacts normal physiological aging, WT mice were treated with stavudine from weaning age and tissue type I interferon expression was assayed at 24 months. We found that treatment of WT aged mice with stavudine significantly reduced type I interferons in several tissues, similar to what was observed in SIRT6 KO animals (**Figure 7D, E**). Stavudine treatment also showed a trend towards reducing plasma concentration of multiple cytokines and chemokines, including those belonging to senescence-associated secretory phenotype (SASP), e.g., IL-6, IGFBP-3, IGFBB-6, CXCL1, CXCL4, TNF RI, CCL5, G-CSF, MCP-1 as measured by antibody arrays in WT aged mice (**Figure S7**). Inducibility of these cytokines by poly(I:C), the inducer of interferon response, remained the same with and without stavudine treatment suggesting that stavudine does not affect interferon signaling pathway per se but rather reduces the levels of endogenous inducers such as cytoplasmic L1 cDNAs.

**Figure 7.**
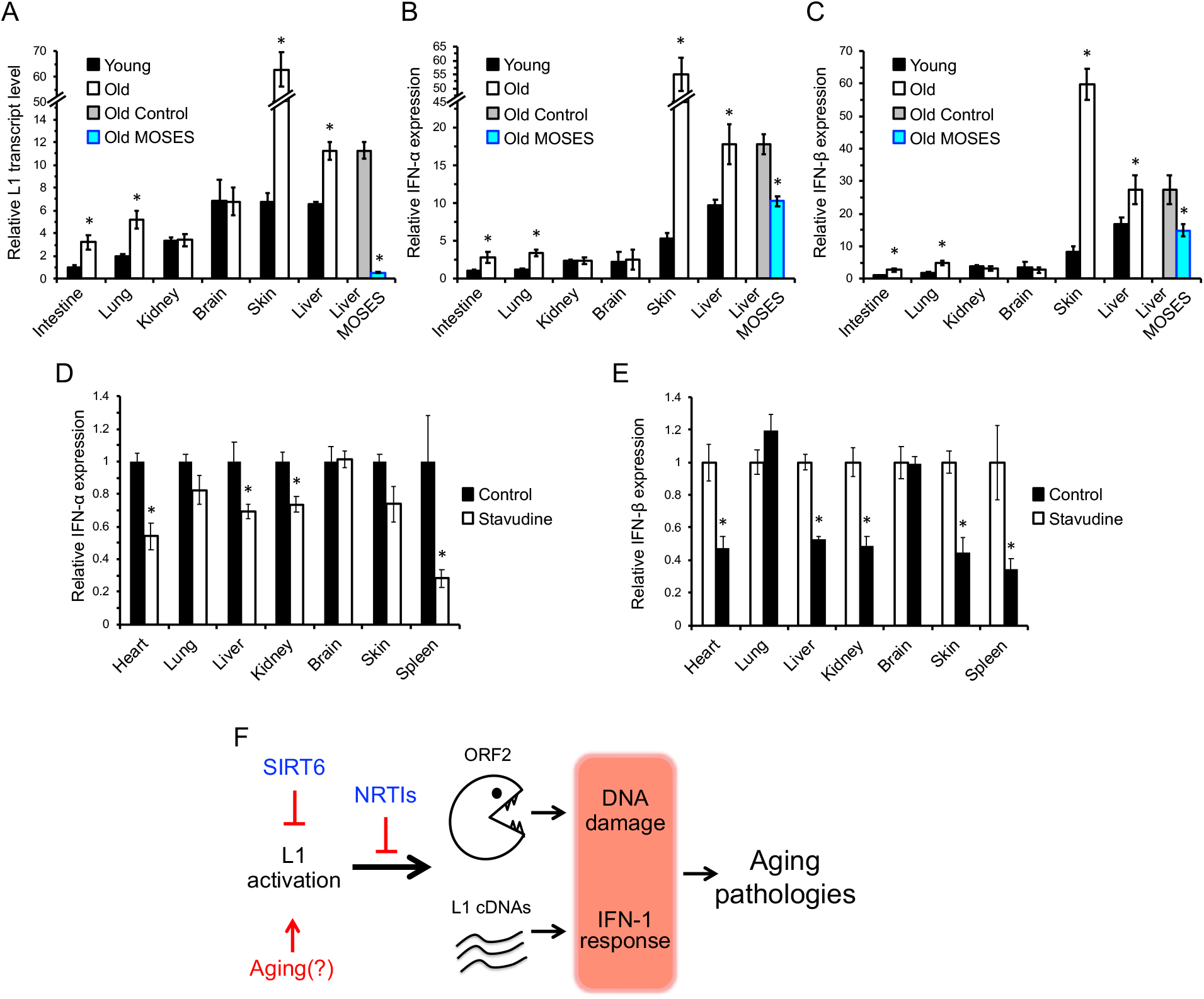
L1 transcripts and interferon response are elevated in WT aged mice and are rescued by treatment with NRTIs and SIRT6 overexpression. **A,** L1 transcripts are elevated in tissues of aged (24 month old) animals, and are repressed in aged MOSES mice overexpressing SIRT6. **B, C,** Aged tissues show elevated IFN-α/β expression whereas MOSES mice show decreased expression. Tissues from 4 months and 24 months old BL/6 and 28 months old MOSES animals were analyzed for L1 expression. Aged MOSES mice have reduced L1 transcripts. Young intestine was used as a reference for the other samples after normalization to actin. Statistical significance was determined by the Student’s *t*-test asterisk indicate *p* < 0.05. **D, E**, Treatment with NRTI reduces IFN-α/β in WT aged mice. **F,** Model. L1 activation causes age-related pathologies through induction of DNA damage and a type I interferon response. These pathologies are rescued or attenuated by inhibition of L1 reverse transcriptase with NRTIs.

To test whether SIRT6 can suppress L1 in the wild type aged mice, we tested RNA from the livers of aged mice constitutively overexpressing SIRT6 (MOSES mice) (Kanfi et al., 2012). Remarkably, the age-related increase in L1 and type I interferon transcription were rescued in aged MOSES mice compared to their littermate controls (**Figure 7A-C**) indicating that elevated SIRT6 activity prevents age-related L1 activation during normal aging. Taken together, these data indicate that L1 suppression not only impacts the progeroid phenotypes observed in SIRT6 KO animals, but similarly impacts normal aging pathologies.

## DISCUSSION

L1s are elevated and, in some cases, causative in pathologies, including inflammation, cancer and neurodegenerative diseases. Here, we demonstrate an active role for these retrotransposons in the progeroid phenotypes associated with SIRT6 deficiency in mice. In this study, we demonstrate that inhibiting L1 RT activity not only alleviates the cellular and physiological dysfunctions of SIRT6 KO mice, it also extends the lifespan of these animals and correlates with observations in WT aged animals. Specifically, we show that L1 activity results in increased cytoplasmic L1 DNA, inducing a type I interferon response through cGAS cytosolic DNA sensing and promoting pathological inflammation. These results, combined with similar observations in aged, WT mice, strongly support a detrimental role for L1s in the process of normal aging. Moreover, this study indicates that L1 activity contributes to the pathology of SIRT6 deficient mice.

It should be noted that NRTIs are not specific to specific reverse transcriptases. There remains the possibility that other retroelements, such as endogenous retroviruses, may contribute to aging pathologies. However, we were able to strongly recapitulate the results in cell culture experiments using NRTIs by using RNAi targeting several conserved regions on L1 families, indicating that L1 activity is the major contributor to the observed pathologies. Indeed, with L1s constituting the most abundant class of retroelements, it is reasonable to surmise that they make the major contribution to the RT-related type I interferon response.

Interestingly, we found that WT aged mouse tissues, similarly to the progeroid SIRT6 KO, display elevated L1 expression and induction of type I interferons. These observations compliment those reported for the higher incidence of DNA damage in aged tissues, similar to that seen in SIRT6 deficient models. Taken together these results suggest that activation of L1 elements contributes to both age-related genomic instability and sterile inflammation associated with aging. The progressive activation of L1s with age may occur due to loss of silencing (De Cecco et al., 2013a; Villeponteau, 1997) and redistribution of chromatin modifiers such as SIRT1 and SIRT6 (Oberdoerffer et al., 2008; Van Meter et al., 2014).

NRTI treatment was able to greatly improve the health and cellular pathologies in SIRT6 KO mice, however it did not fully rescue the shortened lifespan. This indicates that there are other factors outside of L1 activity impacting these pathologies such as misregulation of glucose homeostasis (Mostoslavsky et al., 2006; Zhong et al., 2010). Furthermore, NRTIs rescue the accumulation of cytoplasmic L1s and type I interferon response but do not target L1 transcription which may also be contributing to the burden these abundant retroelements impose on the cell.

NRTIs, themselves, can pose adverse health effects, as patients treated chronically with NRTIs have demonstrated hepaptoxicity due to inhibition of mitochondrial DNA polymerase gamma (Montessori et al., 2003; Wu et al., 2017). Consistent with these reports, we did not observe an improvement in inflammation markers in the liver tissue, save for type I interferons. Additionally, NRTIs are known to inhibit telomerase and are believed to contribute to observations of accelerated aging in HIV patients (Hukezalie et al., 2012; Leeansyah et al., 2013). While NRTI treatment may prove an effective treatment to conditions predicated by LINE1 activity, it would not be an ideal solution to combat normal, physiological aging. As such, the development of more targeted, anti-LINE1 interventions is now a necessary extension of these and other recent findings linking aberrant LINE1 activation to human disease.

Several studies have demonstrated that cytosolic DNAs can trigger innate immune responses that contribute to cellular senescence and aging (Dou et al., 2017)(Gluck et al., 2017)(O’Driscoll, 2008; Thomas et al., 2017; Yang et al., 2017). The identity of the cytosolic DNAs however, has been a mystery. Recently, it has been reported that accumulation of L1 cDNA in the cytosol of TREX1 deficient neurons drives type I interferon production and leads to apoptosis (Thomas et al., 2017). Here we propose that the L1 activation and the resulting cytosolic L1 cDNAs, along with L1-mediated genomic instability contribute to age-related inflammation and other aging pathologies (**Figure 7D**). Thus, L1 inhibition may be a viable strategy for treatment of age-related diseases. Interventions, such as NRTIs, may hold the potential to supplement other treatments or serve as a new form of therapy for age-related pathologies. Future work on effective dosages, late life animal treatments, and developing specific and less toxic anti-L1 inhibitors will pave the way for the future translational applications.

## ACKNOWLEDGEMENTS

This work was supported by grants from the US National Institutes of Health to S.L.H., J.M.S., A.S., A.V.G. and V.G., and Life Extension Foundation to A.S. and V.G.

## AUTHOR CONTRIBUTIONS

MS, AS, and VG designed research and wrote manuscript with input from all authors. MS performed all experiments related to L1 activity, DNA damage, and mouse phenotypic characterization. MVM performed initial experiments described in Figure 1; JA performed mouse NRT treatment and mouse pathology analysis; ZK performed analysis of stem cells; RSG performed pathology evaluation of tissues; TT synthesized Methyl Stavudine; MDC contributed to qRTPCR analysis; KIL performed analysis of cytokine arrays; VK and NN contributed to bioinformatics analysis; SLH contributed to data analysis; AR and HYC provided tissues from MOSES mice; MA, AG, and JMS performed NRTI treatment of aged mice, analysis of cytokines (MA and AG), and contributed to discussion, manuscript writing and data analysis.

## METHODS

### EXPERIMENTAL MODEL AND SUBJECT DETAILS

#### Mice and cell lines

SIRT6 KO and WT mice were acquired from Jackson laboratories (strain 129 Sirt6tm1Fwa/J, 006050) and maintained in accordance with the regulations designated and approved by the University of Rochester’s Committee on Use and Care of Animals. All mouse-derived cell lines were isolated from these animals. The cell lines were established from the mice in the course of this study. The identity of the cell lines was authenticated by PCR genotyping. MOSES mice (Kanfi et al., 2012) were from Bar Ilan University.

### METHODS DETAILS

#### NRTI treatments

Pregnant SIRT6 heterozygous mice were administered 1.5 mg/ml stavudine or lamivudine in drinking water starting immediately after mating. Postnatally, the pups were given 400 mg/kg/day stavudine, or 600 mg/kg/day lamivudine, or water (control) by a flexible pipette into the animal’s mouth once a day.

For studies of WT aged animal, C57BL/6j mice were provided 1 mg/ml stavudine in drinking water for 24 months post-weaning and tissues were harvested at 25 months.

#### Analysis of cytokines in mouse plasma following stavudine treatment

55-week-old C57BL/6j mice were maintained either with or without 1 mg/ml stavudine in drinking water (neutral pH) since weaning (4-5 week old). After reaching 61 weeks of age, mice from each group were either treated or untreated with i.p.-injected poly(I:C), 25 mg/mouse, to induce interferon type I response. Control groups were injected with the vehicle (PBS). Blood was collected 6 hours post treatment using heparin tubes. Equal volumes of plasma were combined from two males and two females from each group for the detection of circulating cytokines and chemokines using Mouse Cytokine Antibody Array C3 kit (RayBiotech, Norcross, GA) according to the manufacturer’s protocol. Briefly, the membranes precoated with capture antibodies were blocked and then incubated with plasma overnight at 4°C (plasma was diluted two-fold with blocking buffer). The membranes were then washed with washing buffer and incubated with biotinylated detection antibody cocktail overnight at 4oC. Following this step, the membranes were washed once again and incubated with streptavidin-HRP for two hours and developed using detection buffers provided in the kit. The immunoblot images were captured and visualized using the BioRad Molecular Imager GelDoc and the intensity of each spot was analyzed using ImageJ software.

#### Cell culture

All cell lines were maintained in humidified incubators at 5% CO_2_, 5% O_2_, at 37°C. Cells were grown in Eagle’s minimum essential medium with 15% fetal bovine serum and 1x penicillin/streptomycin. Drug treated cells were supplemented with 10 μM Stavudine or Lamivudine. The cell lines are routinely tested for mycoplasma contamination.

#### Transfections

Transfections were carried out by plating cells at a density of 500,000 cells/10 cm plate two days prior to transfection. Transfections were carried out using the Amaxa Nucleofector with Normal Human Dermal Fibroblast transfection solution.

#### Quantitative RT-PCR

Total RNA was isolated from cells at 80% confluence using Trizol Reagent and then treated with DNase. cDNA was synthesized using Superscript III (Life Technologies) cDNA kit with the Random Hexamer primer. qRT-PCR was performed on the BioRad CFX Connect Real Time machine with SYBR Green Master Mix (BioRad) using 30 ng of cDNA per reaction with 4x reactions/sample. Efficiency verified mL1 primers previously described were used to assess L1 transcript abundance standardized to QuantumRNA Actin Universal primers (Van Meter et al., 2014). All primers can be found in Key Resources Table.

#### L1 DNA Content

Tissues were harvested from animals and immediately frozen in liquid nitrogen. Tissues were cold processed with pestle and mortar and then genomic DNA was isolated using Quiagen’s DNeasy Blood and Tissue kit. DNA concentration was measured in quadruplicate and then serially diluted to 3 picograms/μl. Two μl of diluted genomic DNA was then loaded into SYBR Green Master Mix reaction and assayed using mL1 primers on BioRad CFX Connect Real Time machine.

#### Cytoplasmic DNA Extraction

For MEFs, cells were grown to 75% confluence, gently trypsinized from the plate, counted and collected by centrifugation. A cytoplasmic lysis solution (Shen et al., 2015) was used to resuspend cells, which were incubated at 4°C on a rotor for 10 min. Nuclei were removed by centrifugation and the supernatant was treated as previously described (Shen et al., 2015). A solution of 3M CsCl, 1M potassium acetate, and 0.67M acetic acid was added to supernatant, incubated on ice for 15 min, then centrifuged to remove any residual cellular debris and genomic DNA before column purification using Qiaquick PCR cleanup columns. Final elutions were quantified and assayed for nuclear genomic contamination using primers for GAPDH and 5S ribosomal subunit with 10 ng of DNA via qPCR.

For organs, organs were harvested from animals and mixed cell-type cultures were generated from the organs in F12 media (Gibco). Cells were passaged once after initial colonies formed from processed tissue. At 75% confluence, cyptoplasmic fractions were isolated by direct application of cytoplasmic lysis solution direction to the plate and incubated at 4°C with gentle shaking for 10 min. Lysates were then collected and processed in the same manner as described above.

#### Mouse activity assays

For flight response assays, animals were placed in an open arena in the center of a 4” radius stage and then timed until they fully exited the stage. Full exit is defined as all four limbs outside of the circle. Each animal was tested 3 times, with 30 min rest intervals in normal housing between trials.

Foraging and exploration activity were assayed by placing animals in a gridded arena for 30 sec at a time. A camera was used to record animal movement, which was then quantified using Tracker software. Animals were allowed to rest in 30 min intervals between trials in normal housing to prevent acclimation to the arena.

Mouse strength was assayed by inverted screen test as described in (Deacon, 2013). A mouse was placed in the center of the wire mesh screen, the screen was rotated to an inverted position over 2 sec, with the mouse’s head declining first. The screen was kept above a padded surface. The time when the mouse fell off was recorded. The following scoring was used: Falling between 1-10 sec = 1; Falling between 11-25 sec = 2; Falling between 26-60 sec = 3; Falling after 60 sec = 4. For each animal the point score was calculated as a sum of the scores for three trials.

#### Immunofluorescence and Apoptosis

γH2AX and 53PB immunostaining was carried out as previously described (Mao et al., 2011). Anti- γH2AX and anti-53BP1 antibody was purchased from Abcam (ab22551 and ab36823). For ssDNA immunostaining, cells were fixed with PFA, followed by 24hr methanol incubation at -20°C. Cells were then incubated at 37°C for 4hrs with RNaseA. Antibody from Millipore (MAB3868). Apoptosis in fibroblasts was measured using the Annexin V Staining Kit (Roche). Apoptosis in tissues was measured using TUNEL method with *in situ* apoptosis detection kit, Abcam (ab206386).

#### Histology

Mouse tissue was fixed in Bouin’s fixative prior to being embedded in paraffin. Fixed tissues were then sectioned at 6 μm and hematoxylin/eosin staining was performed by standard methods at the University of Rochester Medical Center’s HBMI core.

#### Synthesis of 5’-*O*-Methyl stavudine

Reaction was conducted in oven-dried glassware under an argon atmosphere. Reaction solvents were obtained from a solvent system by Innovative Technologies Inc. Reagent grade solvents were used for all work-up procedures and extractions. Stavudine was purchased from Matrix-Scientific, sodium hydride (60 % dispersion in mineral oil) and methyl iodide were purchased from Sigma-Aldrich. Nuclear magnetic resonance spectra were obtained on a Bruker Avance 500 MHz spectrometer. Mass spectra were obtained using a Shimadzu LCMS-2010 mass spectrometer.

To a solution of stavudine (224 mg, 1 mmol) in dry THF/DMF (6 mL, 5:1 v/v) was added sodium hydride (400 mg, 10 mmol). The reaction mixture was stirred for 10 min under an argon atmosphere. The reaction mixture was then cooled to 0°C, and methyl iodide (67 μL, 1 mmol) was added slowly. The reaction mixture was stirred overnight while temperature was slowly raised from 0°C to room temperature. The reaction was quenched by methanol, neutralized by acetic acid, then volatiles were removed by evaporation. DMF was removed by azeotropic distillation with toluene. The resulting solid was suspended in DCM, and the mixture was washed with aq. NaHSO_3_. The organic layer was dried over magnesium sulfate, concentrated *in vacuo*, and then purified by silica gel chromatography (elution by 1:1 then 1:4 hexanes/EtOAc) to afford 148 mg of the final product, > 97% purity. The spectroscopic and spectrometric data were consistent with those reported in the literature (Fowler et al., 2014).

#### Comet assays

Mouse embryonic fibroblasts were maintained in control or drug treated media for 15 population doublings and then analyzed for DNA damage using the Trevigen CometAssay system. In brief, non-confluent cells were embedded in agarose, fixed, and then subjected to electrophoresis in a neutral buffer solution. Cells were then stained using SYBR Gold and analyzed by microscopy. Quantitative analysis was performed using CASP software.

#### Blood glucose analysis

Blood was collected from tail bleeds at the indicated age, and serum glucose was measured. Serum glucose was measured using One-Touch Ultra-2 blood glucose glucometer kit, per the manufacturer’s instructions.

#### Bone Marrow Stem Cell Counting

Bone marrow nucleated cells were isolated from 2 femurs and 2 tibias from each mouse, stained with trypan blue and counted. Two million bone marrow cells were stained with DAPI and antibodies (BioLegend): anti-mouse lineage-Pacific blue, anti-mouse Sca1-APC, anti-mouse c-Kit-PE/Cy7, anti-mouse CD48-APC/Cy7, anti-mouse CD150-PerCP/Cy5.5, followed by flow cytometry analysis. In methylcellulose assay, 50,000 cells per ml were plated onto ultra-low attachment 35mm plates, incubated in a humidified 37°C incubator with 5% CO_2_ and 20% O_2_. Colonies were scored 12-14 days later.

#### LINE1 RNAi Vector Systems

The ABM Good iLenti siRNA vector system was used to generate the siRNA cassette using the conserved L1 sequence, TGGACCAGAAAAGAAATTCCTC.

The **BLOCK-iT Pol II** miR RNAi Expression Vector was used to generate the shRNA expressing vector. Five shRNAs were combined into the final vector. The targeting sequences are as follows; shRNA 1: TCCAAATAGACTGGACCAGAA, shRNA 2: AGAGCCTGGACAGATGTTATA, shRNA 3: GAGGAGTAGACGGCAGGAAAT, shRNA 4: TGGGATTAGTGCAGAGTTCTA, shRNA5: CCATACTTATCTCCTTGTACT

#### Immunoprecipitation of cGAS

Cultured MEFs were washed with PBS prior to crosslinking and fixation. 200mJ of UV radiation was applied using a Stratolinker UV system, followed by cell harvest and counting. Cells were then fixed for 10 min in 4% PFA and washed 3x with PBS. Cells were then lysed for 30 min at 4 °C while rotating using a lysis buffer containing 20 mM Tris-HCl [pH 7.5], 0.5 mM EDTA, 150 mM NaCl, 10 mM KCl, 0.5% Triton, 1.5 mM MgCl2, 10% glycerol, 0.2 mM PMSF, 10 mM β-mercaptoethanol. Lysates were cleared by centrifugation at 12,000g for 30 min at 4 °C. cGAS was bound using either 5ug cGAS D3080 (Cell Signaling) or ABF124 (Millipore) for an overnight incubation, followed by the addition of 30 μl of Agarose A beads with salmon sperm DNA and 2h incubation while rotating. Beads were washed 5x with lysis buffer before DNA isolation using Qiagen DNeasy Blood and Tissue kit.

### QUANTIFICATION AND STATISTICAL ANALYSIS

Unless otherwise noted, the Student’s t-test was used to determine statistical significance between groups. Lifespan assays were analyzed with the log-rank test. Body weight curves were analyzed using one-way ANOVA. All tests were two-tailed and p-values were considered significant below a 0.05 threshold. Cell culture experiments utilized at least two separately derived cell lines for each genotype and were performed in triplicate unless noted otherwise. For animal studies the sample size reflects a balance between the minimum numbers necessary for efficient analysis and some surplus animals to take into account individual variation.

### KEY RESOURCES TABLE

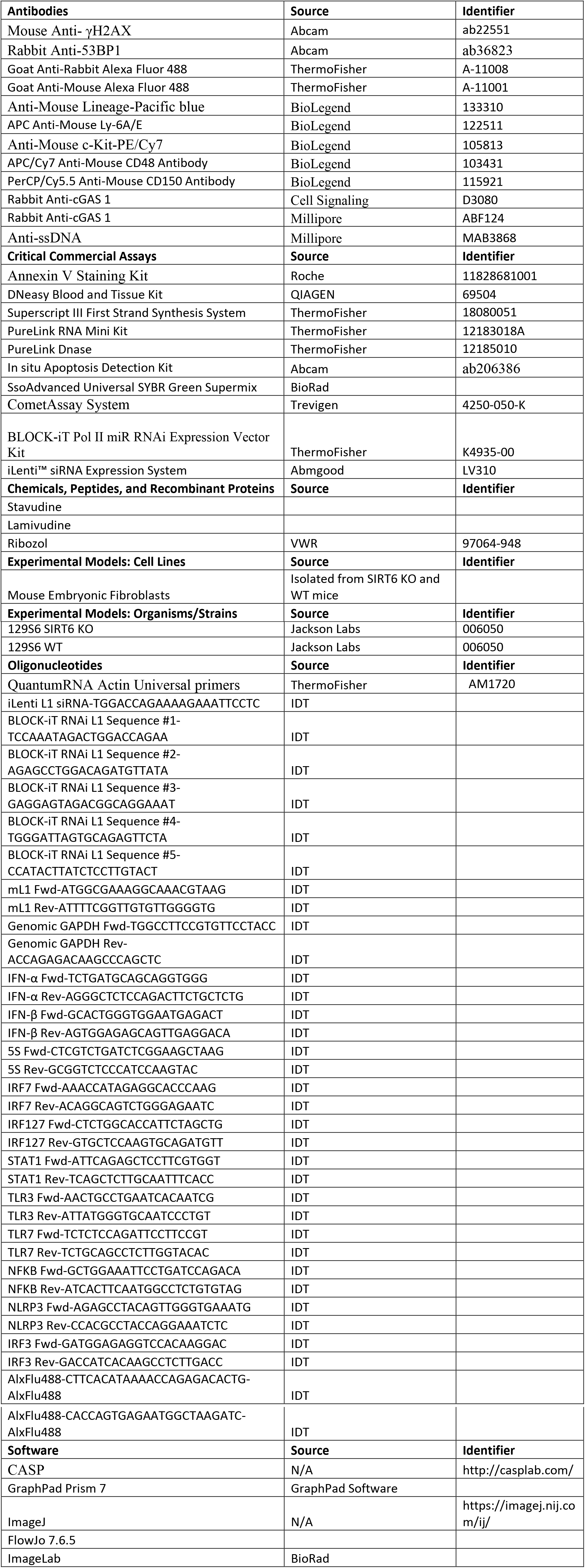

## SUPPLEMENTARY VIDEOS

Mice perform in open field and foraging activity tests. Representative videos of WT, SIRT6 KO control, SIRT6 KO treated with lamivudine, and SIRT6 KO treated with stavudine are shown. Mice were tested at 27 days of age.

## Supplemental Information

- Figures S1-S7

- Supplemental videos 1-4, showing WT and SIRT6 KO mice treated or not with stavudine and lamivudine performing in open field tests.

**Figure S1. Related to Figure 1.**
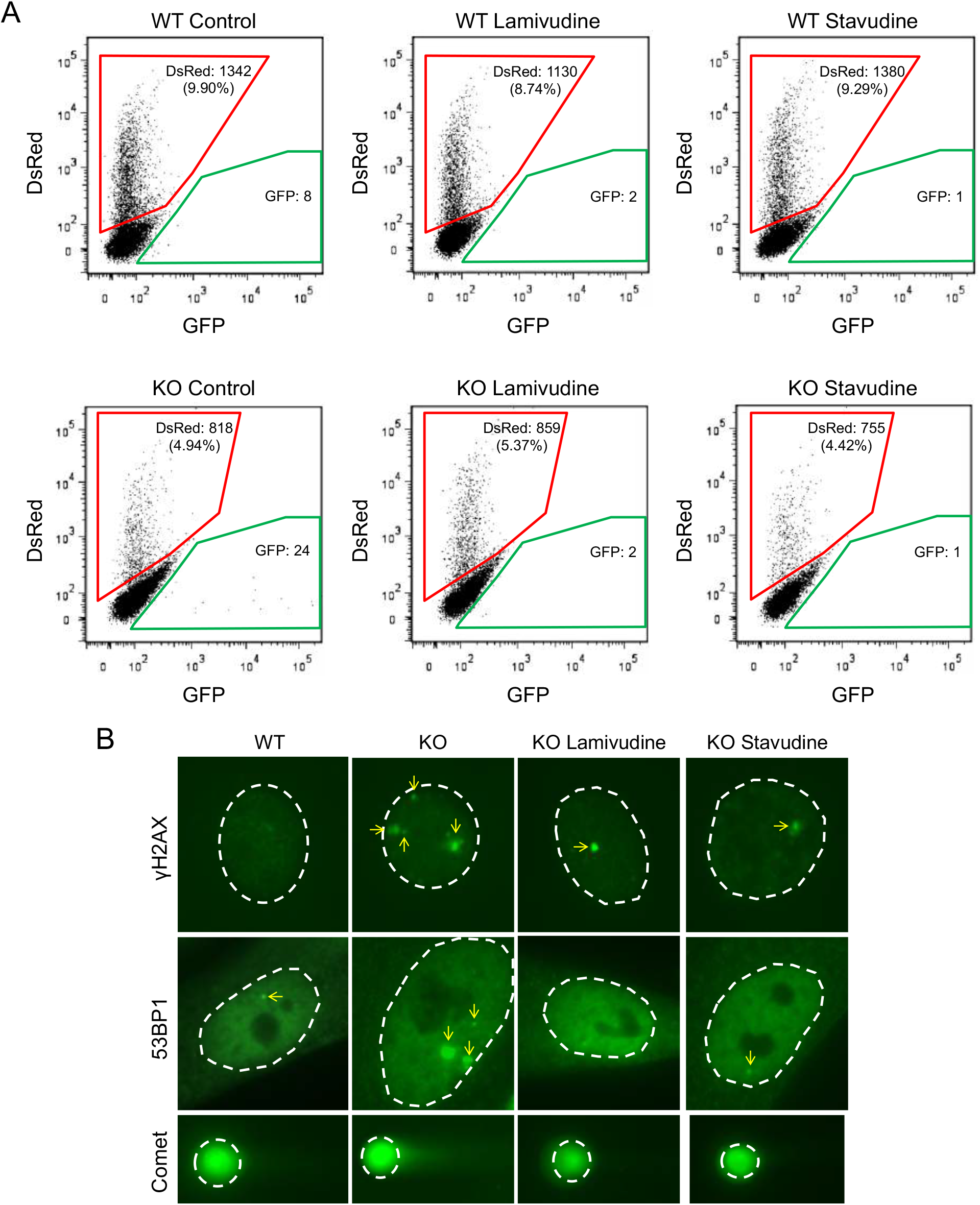
Representative FACS traces for L1 EGFP reporter and representative images for γH2AX, 53BP1 immunostaining and comet assay. **A,** Representative FACS traces from WT and SIRT6 KO MEFs transfected with mL1-EGFP reporter plasmid. A transfection control plasmid containing dsRed was used to standardize the retrotransposition activity with transfection efficiency. **B,** Representative images for γH2AX, 53BP1 immunostaining and comet assay. Dashed white lines indicate the nucleus of the cells. Yellow arrows indicate foci for 53BP1 and γH2AX.

**Figure S2. Related to Figure 4.**
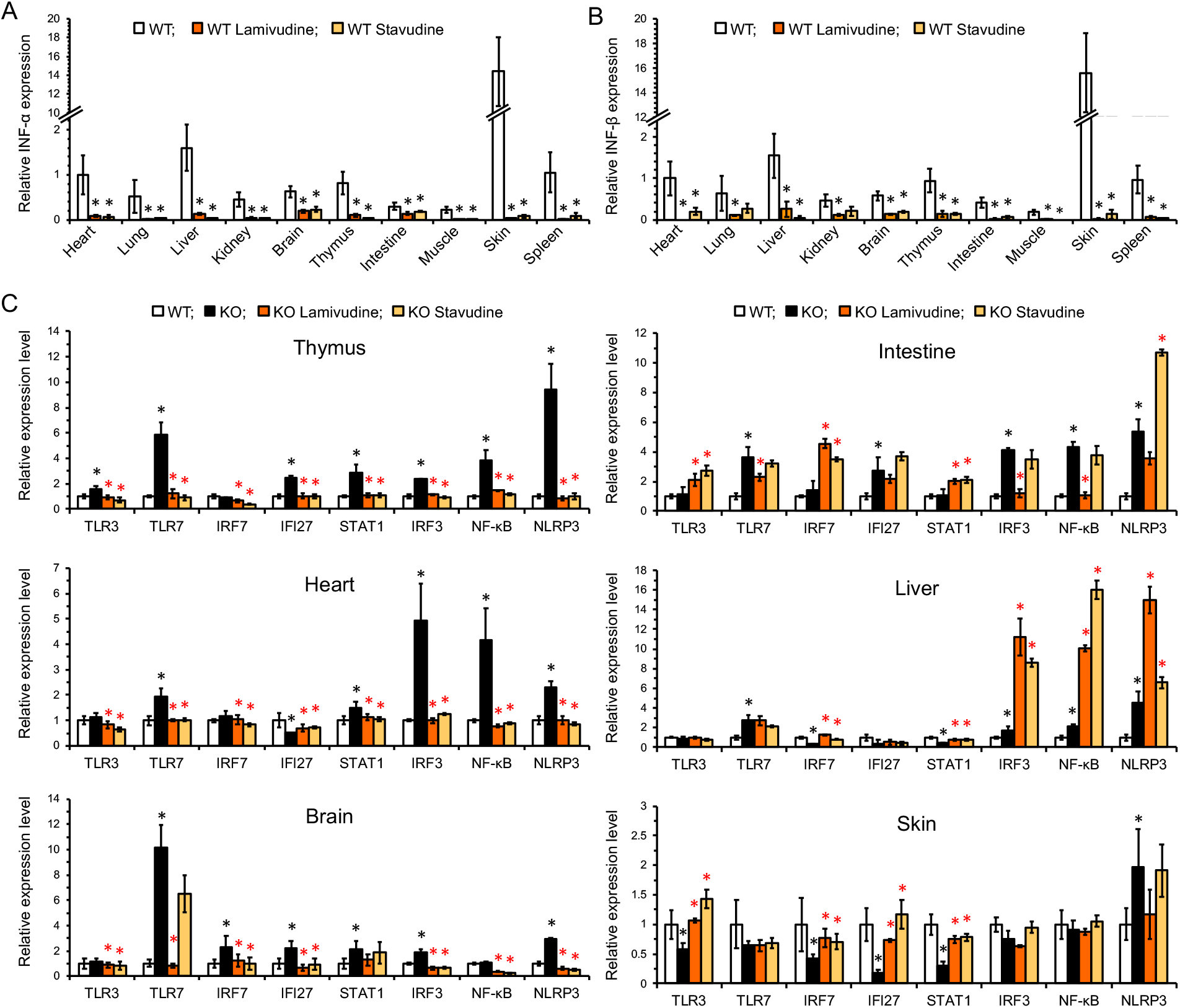
Expression of inflammation markers in tissues of mice treated with NRTIs. **A, B,** NRTI treatment reduces basal levels of IFN-α/β in WT animals. The qRT-PCR values were normalized to actin and WT heart was used as a reference. Three animals were assayed for each group and error bars show s.d. Statistical significance was determined by the Student’s *t*- test, asterisk indicates *p* < 0.05. **C,** RT-qPCR analysis of inflammation genes in major organ systems was carried out on 27-day old animals. All values are relative expression levels compared to actin and then normalized to the WT. Data represents averages from five animals per treatment. Error bars show s.d. Statistical significance was determined by the Student’s *t*-test. Black asterisks indicate significant differences from WT, and red asterisks indicate significant differences from water-treated SIRT6 KO, *p* < 0.05.

**Figure S3. Related to Figure 5.**
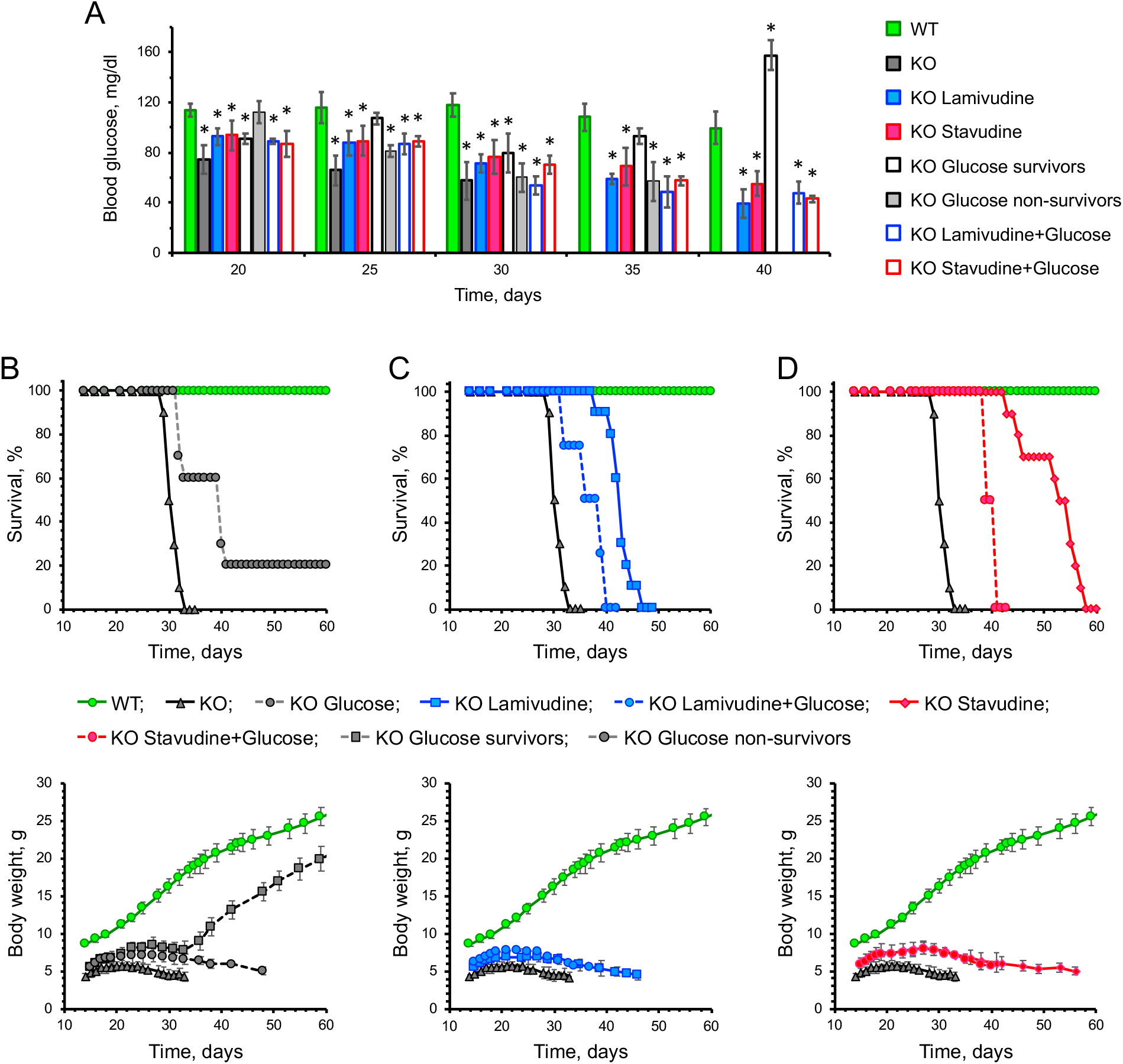
Glucose supplementation does not enhance NRTI impact on blood glucose, lifespan or bodyweight. **A,** Blood glucose levels in NRTI-treated animals. NRTI treated animals still had low blood glucose at the time of weaning (day 22) compared to the WT, but slightly delayed further decline in blood glucose at day 30. Supplementation of 10% glucose in both control and NRTI treated animals did not significantly impact blood glucose levels. Five animals were assayed for each group and error bars indicate s.d. Asterisks indicate values significantly different from WT. Significance was determined by the paired *t*-test, p < 0.05. **B-D,** Glucose supplementation does not improve NRTI rescue of SIRT6 KO mice lifespan or bodyweight. Control and NRTI-treated animals had diets supplemented with 10% glucose. A subset of control animals supplemented with glucose showed increased lifespan and bodyweight gain. NRTI-treated animals with glucose supplementation showed a significant decrease in lifespan compared to NRTI-only controls and no change in bodyweight. Ten animals were assayed for each group. Lifespans analyzed by log-rank and body weight by one-way ANOVA, *p* < 0.05.

**Figure S4. Related to Figure 5.**
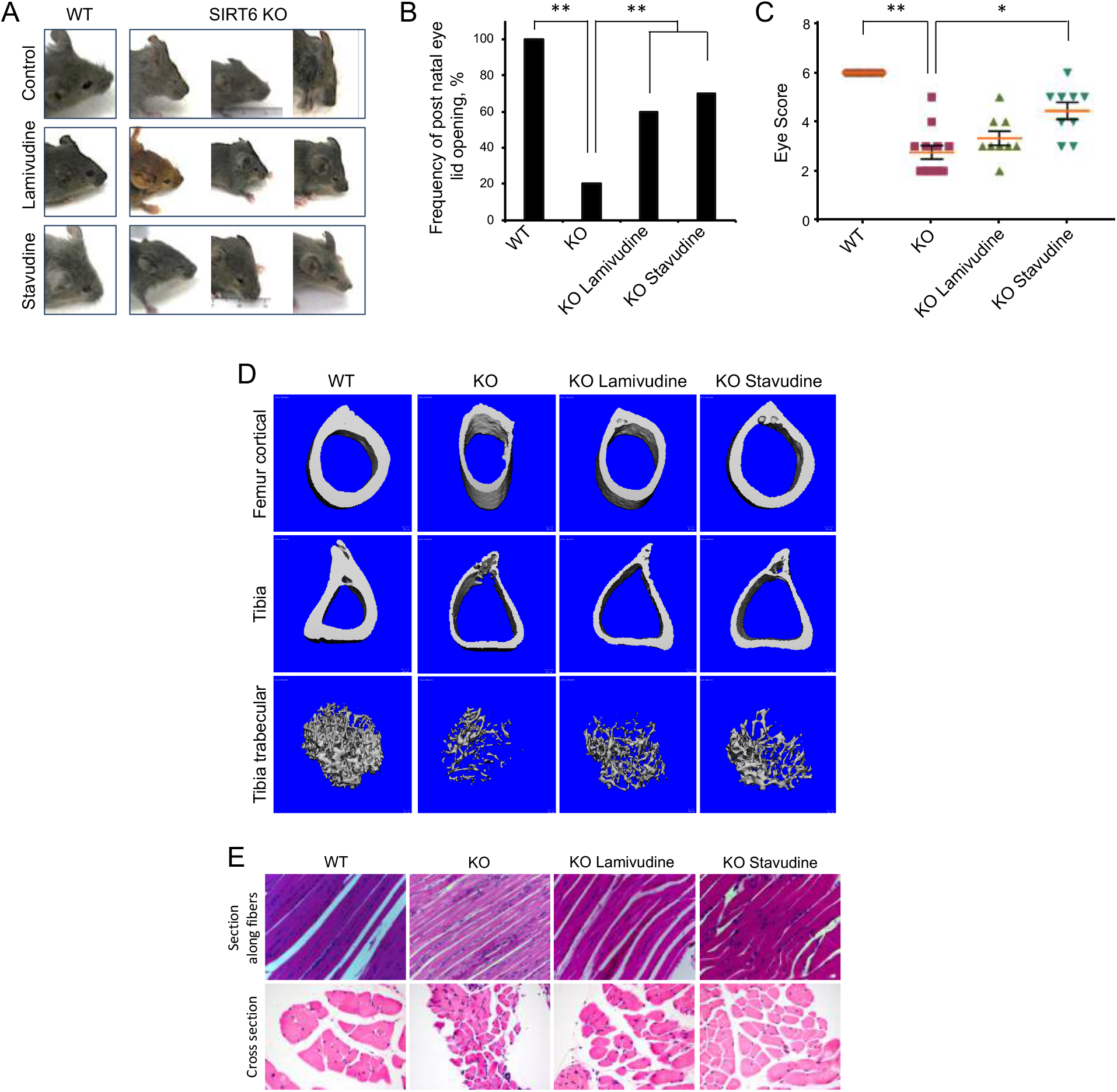
NRTI treatment improves SIRT6 KO physical robustness. **A**, SIRT6 KO pups display a significantly reduced frequency of postnatal eyelid opening and their eyes remain closed until their death. Pups are pictured at 30 days old. Stavudine and lamivudine treatment increased the frequency of eye opening in the SIRT6 KO mice. **B**, Percentage of mice that opened their eyes by day 30. Ten mice were analyzed for each group. Double asterisk indicates *p* < 0.0001, Student t-test. **C**, Individual animal eye scores. 1=fully closed, 2=partially open (some eyeball visible between eyelids), 3=fully open. Ten mice were analyzed for each group. Asterisk indicates *p* = 0.001, double asterisk indicates *p* < 0.0001, Student t-test. **D,** Representative images of femur and tibia cross-sections and tibia trabecular. Images were obtained by CT-scan of analysis of 27-day old animals. **E**, Representative cross sections stained with hematoxylin & eosin of quadriceps of WT or KO NRTI-treated animals.

**Figure S5. Related to Figure 5.**
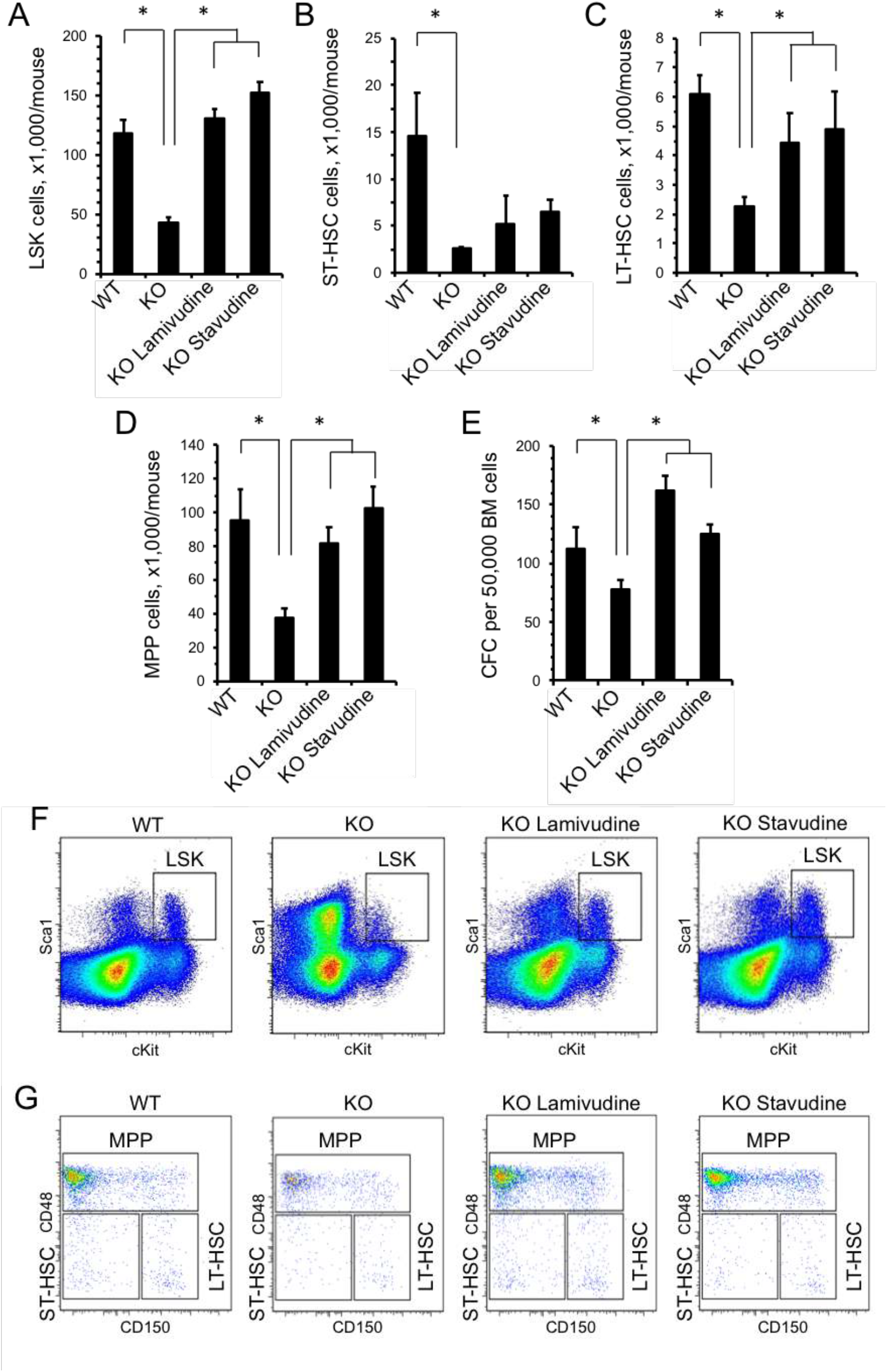
SIRT6 KO mice have reduced bone marrow stem cell populations. **A-D,** The frequency of LSK, LT-HSCs (Lin-Sca1^+^cKit^+^CD48^-^CD150^+^), ST-HSCs (Lin^-^ Sca1^+^cKit^+^CD48^-^CD150-), HSPCs(Lin^-^Sca1^+^cKit^+^), and MPPs(Lin^-^Sca1^+^cKit^+^CD48^+^CD150^+^) in the bone marrow was determined by flow cytometry. Flow cytometry plots are gated on Lin^-^ bone marrow cells. Data presented are the number of specified cell population per mouse. **E,** The number of colonies formed in methylcellulose colony-forming assay using bone marrow cells from WT, SIRT6 KO, and NRTI-treated mice. **F, G**, Representative flow cytometry analysis for Lin-bone marrow cells from WT, SIRT6 KO, and NRTI-treated mice. At least three animals were assayed for each group and error bars show s.d. Statistical significance was determined by the Student’s *t*-test; asterisk indicate *p* < 0.05.

**Figure S6. Related to Figure 5 and Figure 6.**
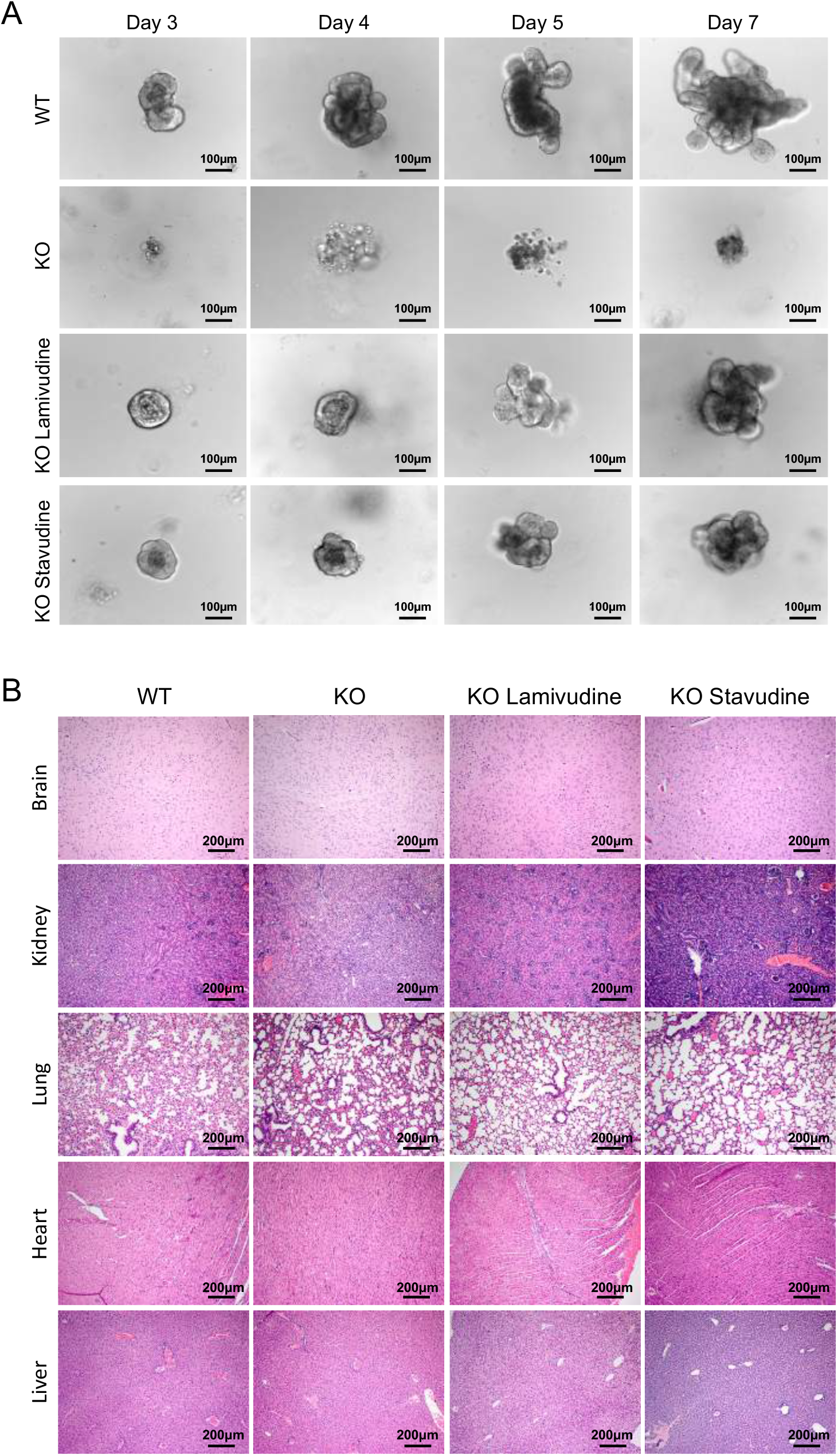
Tissue histology for major organ systems in SIRT6 KO mice following NRTI treatment. **A,** Crypt cells were isolated from 2 cm sections of mouse intestine and cultured in matrigel. WT crypts quickly give rise to organoids and display distinct budding by day 7, whereas SIRT6 KO crypt cells showed little organoid development and often failed to form full organoids. NRTI-treated SIRT6 KO crypts displayed organoid growth and development, but with reduced budding compared to WT organoids. **B**, All animals were euthanized at 27 days of age and tissues were immediately prepared for histology. Most tissues show little difference between WT, SIRT6 KO, and NRTI-treated animals. Images are representative of collected images from two animals per treatment. Tissues are hematoxylin & eosin stained at 100x magnification (scale bars = 200 μm).

**Figure S7. Related to Figure 7.**
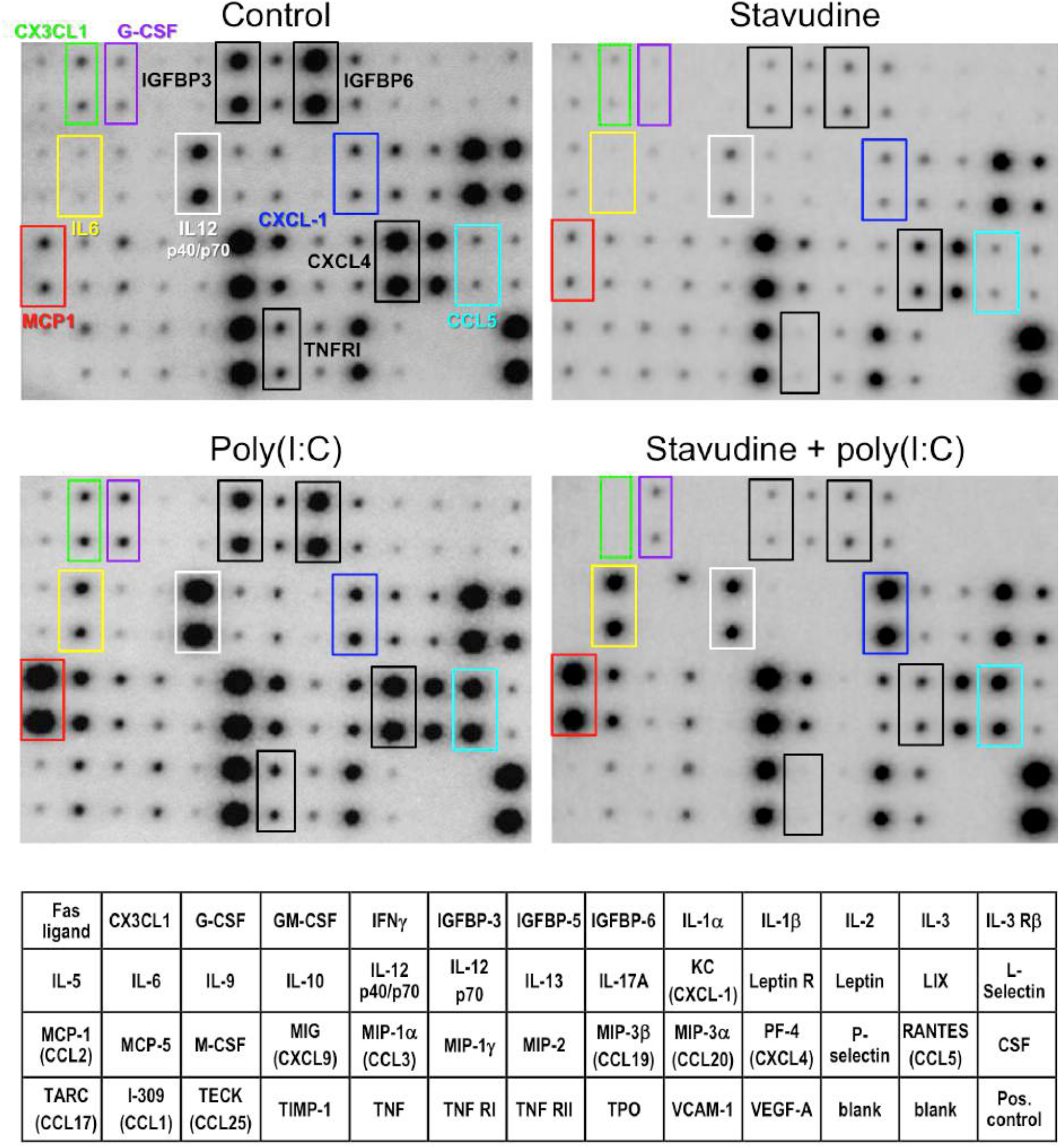
Life-long stavudine treatment reduces concentration of multiple cytokines and chemokines in plasma of aged WT mice. Wild type C57BL/6 mice were treated with 1 mg/ml stavudine in drinking water since weaning. At 61 weeks of age, mice were treated with i.p.-injected poly(I:C), to induce type I interferon response. Control groups were injected with the vehicle (PBS). Blood was collected 6 hours post treatment equal volumes of plasma were combined from two males and two females from each group for the detection of circulating cytokines and chemokines using Mouse Cytokine Antibody Array C3 kit. Factors that do not change their expression in response to poly(I:C) but are downregulated in stavudine-treated mice are framed in black rectangles. A subset of factors induced by poly(I:C) and downregulated by stavudine are framed in colored rectangles.

